# Ecdysone acts through cortex glia to regulate sleep in *Drosophila*

**DOI:** 10.1101/2022.08.24.505204

**Authors:** Yongjun Li, Paula Haynes, Shirley L. Zhang, Zhifeng Yue, Amita Sehgal

## Abstract

Steroid hormones are attractive candidates for transmitting long-range signals to affect behavior. These lipid-soluble molecules derived from dietary cholesterol easily penetrate the brain and act through nuclear hormone receptors (NHRs) that function as transcription factors. To determine the extent to which NHRs affect sleep: wake cycles, we knocked down each of the 18 highly conserved NHRs found in *Drosophila* adults and report that the ecdysone receptor (EcR) and its direct downstream NHR Eip75B (E75) act in glia to regulate the rhythm and amount of sleep. Halloween genes, a set of ecdysone synthesis genes, have little to no expression in the fly brain, while mRNA levels of the ecdysone target E75 cycle in the fly head, suggesting that glial ecdysone comes from the periphery and may enter the brain more at night. Anti-EcR staining localizes to the cortex glia in the brain and functional screening of glial subtypes revealed that EcR functions in adult cortex glia to affect sleep. Cortex glia are implicated in lipid metabolism, which appears to be relevant for actions of ecdysone as ecdysone treatment reduces lipid droplet size in these cells. In addition, sleep-promoting effects of exogenous ecdysone are diminished in *Lsd-2* mutant flies, which are lean and deficient in lipid accumulation. We propose that ecdysone is a systemic secreted factor that modulates sleep by stimulating lipid metabolism in cortex glia.

**Graphical Abstract:** 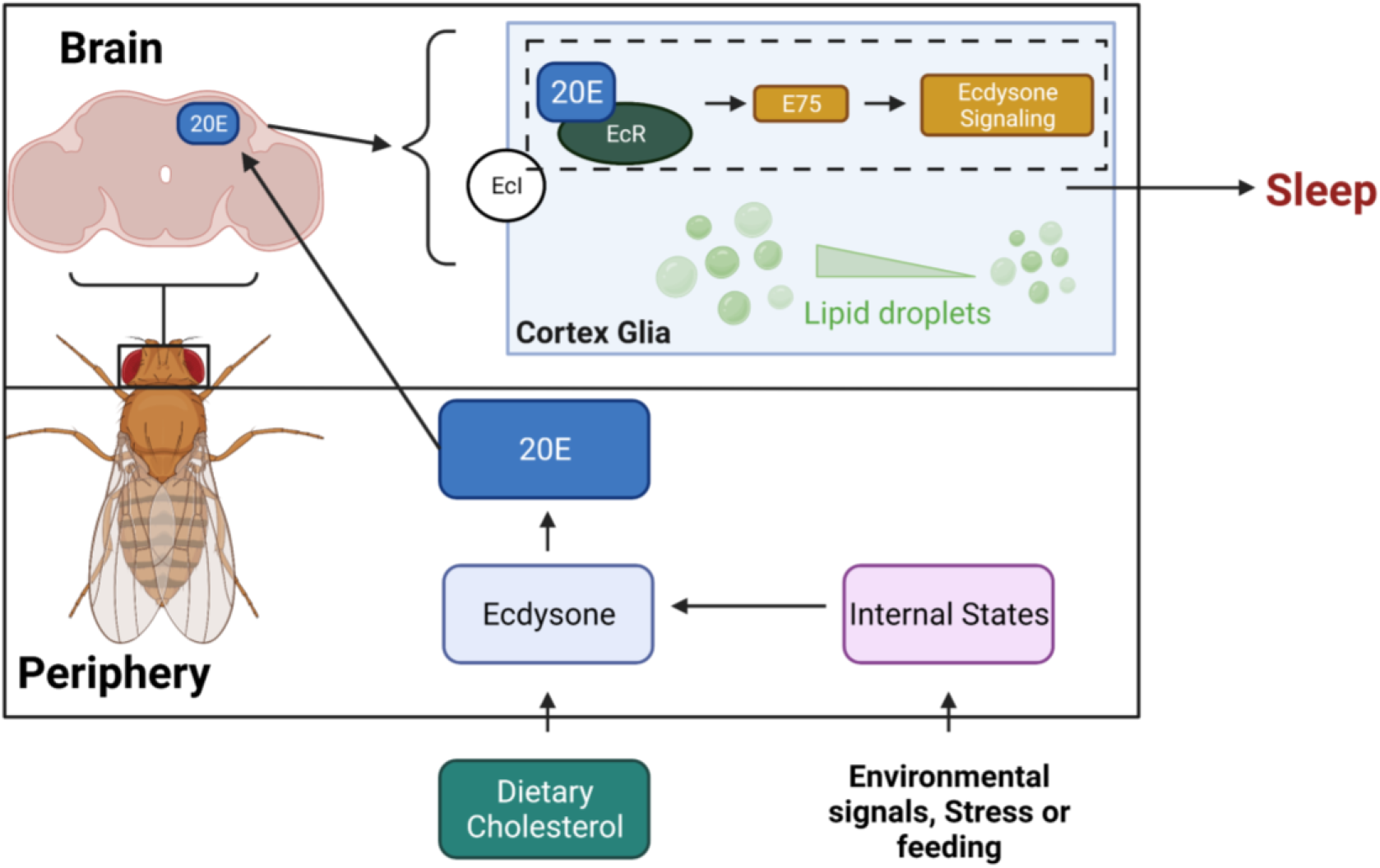

**Highlights:**

- Glial knockdown of ecdysone inducible NHRs reduces sleep.
- The ecdysone receptor (EcR) and its downstream target E75 function in cortex glia to modulate sleep.
- Ecdysone synthesis genes are not expressed in the fly brain, suggesting that glial ecdysone comes from the periphery.
- Ecdysone promotes sleep by mobilizing lipid droplets stored mainly in glia.

## Introduction

Sleep remains a major mystery of biology, with little known about its regulation and function. However, based on ongoing studies, much of them from small animal models, it is becoming apparent that sleep is more than a brain-regulated process that only serves the brain (Borbély et al., 2016; Dubowy and Sehgal, 2017; Harbison et al., 2009; Hobson, 2005; Tononi, 2000). Loss of sleep has systemic effects and is associated with molecular changes in the periphery (Anafi et al., 2013; Chua et al., 2015). In addition, some sleep-promoting effects have been mapped to tissues outside the brain, although the nature of the sleep-regulating signals and their mode of transmission to relevant cells in the brain are poorly understood (Borniger and de Lecea, 2021; Ehlen et al., 2017; Toda et al., 2019).

As a major communication system of the body, endocrine signaling is a promising candidate for mediating long-range effects on sleep (Mong et al., 2011; Morgan and Tsai, 2015; Roller, 2021). Glands of the endocrine system make and release chemical messengers called steroid hormones, which circulate in the blood until they reach their target cells and bind to specific steroid nuclear hormone receptors (NHRs) in the cytosol (Acconcia and Marino, 2016; Zubeldia-Brenner et al., 2016). Given that circulating steroid hormones are lipophilic and can easily enter the brain, together with the broad expression of NHRs in the central nervous system, it is reasonable to infer that NHR signaling is a significant contributor to brain function (Hughes, 2007; King-Jones and Thummel, 2005; Kininis and Kraus, 2008). However, the response mediated by the endocrine system is modulatory, in that it is slower and more long-term than fast, synaptic neurotransmission (Morgan and Tsai, 2015). Thus, the endocrine system drives responses to environmental cues and innate signals critical for development, growth, and metabolism (Kannangara et al., 2021; Oostra et al., 2014). Sleep is a chronic behavior regulated over a relatively long timeframe (a daily cycle), so one might expect it also to be sensitive to endocrine signaling. Indeed, endocrine dysfunction affects sleep, and conversely, sleep and sleep loss affect hormone production and hormonal function, but the specificity and mechanisms underlying these interactions remain to be elucidated (AlDabal and BaHammam, 2011; Kim et al., 2015a, 2015b; Teran-Perez et al., 2012).

To explore the impact of endocrine signaling on sleep, we used a *Drosophila* model and knocked down each of the known *Drosophila* NHRs in neurons and glia. We found that the ecdysone receptor (EcR) and Eip75B (E75) function in both cell types to modulate sleep but have more potent effects in glia. EcR knockdown in glia significantly decreases sleep and disrupts circadian rhythms, as does the knockdown of the newly identified ecdysone importer (EcI). EcR functions in cortex glia to affect sleep, and it appears to do so by regulating lipid metabolism. Together these studies identify a long-range signal that modulates sleep in adult *Drosophila*.

## Results

### Pan-neuronal and pan-glial knockdown of EcR and E75 reduce sleep

To identify NHRs that affect sleep in adult *Drosophila,* we conducted a systematic genetic knockdown screen of all known NHRs. To avoid developmental effects, which are prominent for most NHRs, we sought to address their role in sleep by knocking them down specifically at the adult stage (Nicholson et al., 2008). Based upon single-cell RNAseq data showing that nearly half of known NHRs are highly expressed in adult fly brain both neurons and glia, we used pan-neuronal and pan-glial GeneSwitch drivers induced in adults with RU486 (nSyb-GS and Repo-GS respectively) to knock down individual NHRs (Davie et al., 2018; Nicholson et al., 2008). All NHRs were targeted either by two separate RNAi lines or UAS-miRNA transgenic fly lines that polycistronically express two independent miRNAs in the same line (Lin et al., 2009). Surprisingly, we found that knockdown of most NHRs reduced adult sleep by varying amounts compared with their GS controls and UAS controls (**Figure1B–1C**). Among them, knockdown of Ecdysone receptor (EcR), the receptor of the primary steroid hormone ecdysone, and its direct downstream targeting gene Eip75B (E75) had the greatest effects on total sleep. Because we noticed that Repo-GS>Wcs control flies sometimes have very variable daytime sleep, we also analyzed all flies’ nighttime sleep, which is quite stable in all conditions. Importantly, EcR and E75 knockdown affect primarily nighttime sleep and phenotypes are stronger when they are knocked down in glia.

**Figure1.**
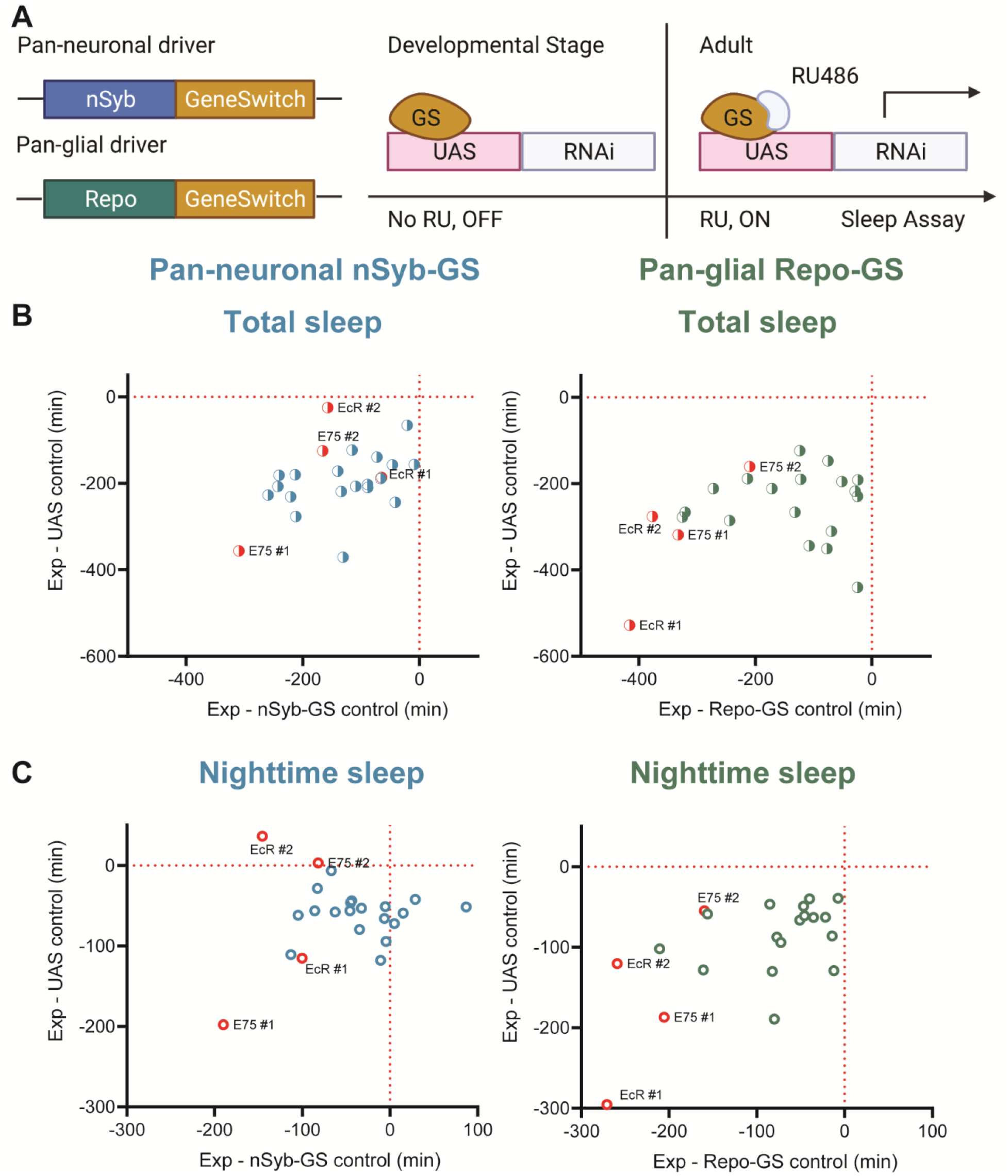
A screen of all nuclear hormone receptors (NHRs) in *Drosophila* identifies sleep-regulating functions of EcR and its downstream target, NHR E75. (A) Flies carrying a pan-neuronal driver nSyb-GS or pan-glial driver Repo-GS were crossed with different UAS lines carrying RNAi constructs against NHRs, and their 5–7-day old F1 female progeny were loaded into DAM monitors to record their activity under 12 hour:12 hour light: dark cycles. GeneSwitch remains inactive during developmental stages and is activated by RU486 in the food in DAM monitors. Behavior data were collected by the DAM system. (B-C) Mean total and nighttime sleep for each group was calculated by Pysolo. Differences between experimental flies and GS and RNAi controls were calculated separately in each independent experiment, and the average values comparing each experimental to its GeneSwitch control (X-axis) and RNAi control (Y-axis) are shown in the plots. The exact numbers of flies used per line are provided in the supplementary Table S1 and Table S2. While knockdown of most NHRs reduces sleep, effects of EcR RNAi #1, EcR RNAi #2, E75 RNAi #1 predominate, especially in glial knockdown experiments.

Across multiple experiments, we found that flies with glial knockdown of EcR slept on average ~470 mins, much less than those with neuronal knockdown that slept ~707 mins. In addition, the total sleep amount of nSyb-GS>EcR RNAi flies was not significantly reduced compared with nSyb-GS control flies, suggesting that glial EcR has a larger role in determining sleep amount (**Figure2A–2D**). Knockdown of E75 in either neurons or glia led to severe sleep loss, with ~419 mins and ~461 mins of daily sleep respectively (**FigureS1A–1D**). Because ecdysone is the most common steroid hormone in *Drosophila,* it directly or indirectly affects multiple NHRs (King-Jones and Thummel, 2005). Indeed, we found that knockdown of ecdysone responsive NHRs reduced sleep more than knockdown of non-ecdysone responsive NHRs, and from the screen of neuronal and glial knockdown, top hits were known Ecdysone relevant NHRs (**TableS1 and Table S2**). If we use 200 min sleep loss compared with both control groups as a cutoff, then 3 out of 4 top hits with neuronal knockdown and 5 out of 7 top hits with glial knockdown are ecdysone relevant NHRs. Genes showing the most dramatic sleep loss when knocked down in glia were EcR, E75, Usp, and Hr39, all directly relevant to the ecdysone signaling pathway. The efficacy of our screen was supported by the identification of non-ecdysone responsive NHR Hr51 *(unfulfilled)* as a positive hit. We found that neuronal, but not glial, knockdown of Hr51 *(unfulfilled),* previously shown to exert circadian effects by acting in central clock neurons (Nagoshi et al., 2012), resulted in an arrhythmic phenotype under 12:12 light: dark conditions (data not shown).

**Figure2.**
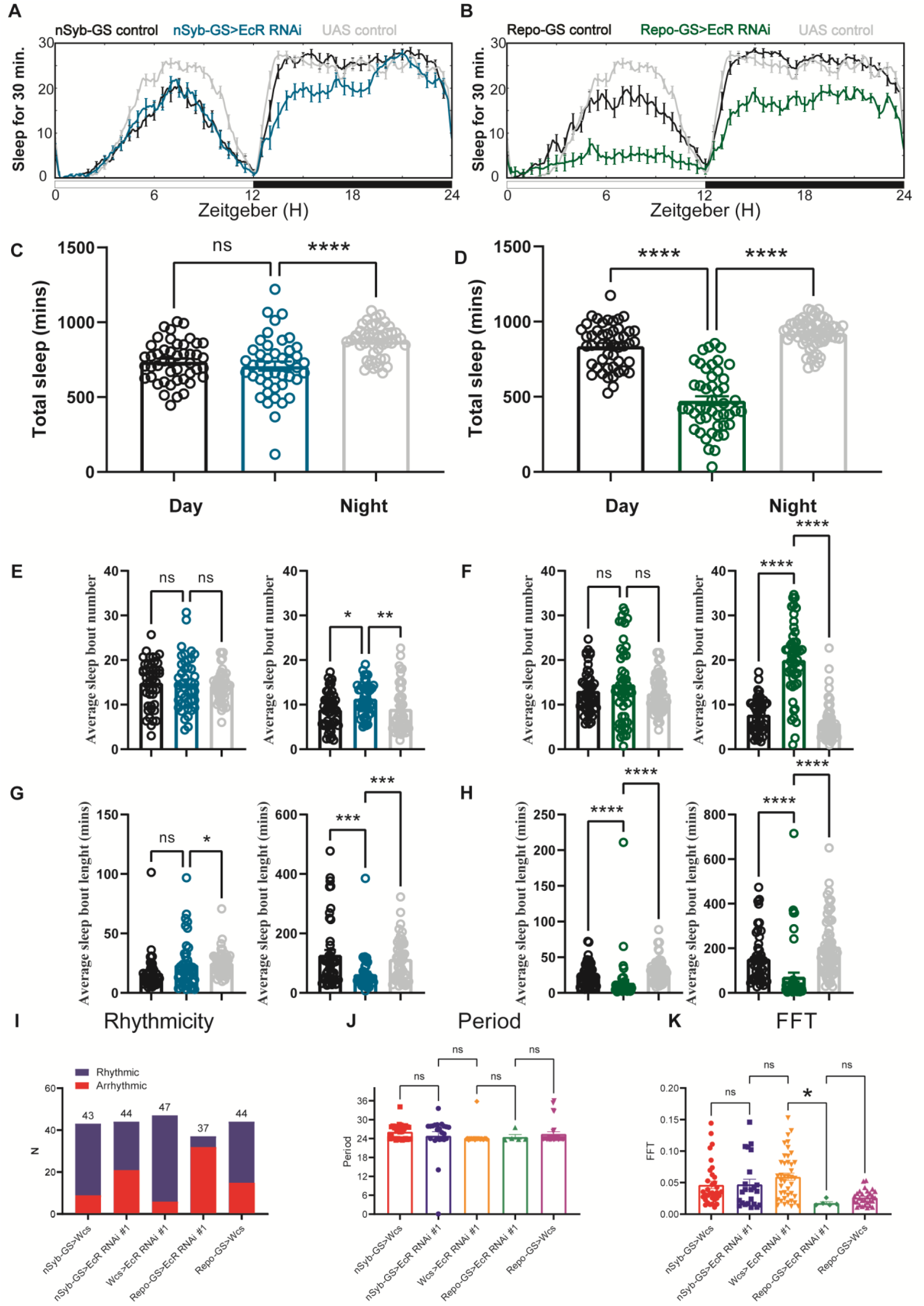
Baseline sleep phenotypes resulting from pan-neuronal or pan-glial knockdown of EcR. (A-B) Show representative sleep traces of nSyb-GS>EcR RNAi #1 and Repo-GS>EcR RNAi #1. N=14-16 per genotype. Data are based on at least three independent experiments. Representative sleep traces are showed because Pysolo we used does not allow combining data from repeated experiments. (C-D) Total sleep of the nSyb-GS>EcR RNAi #1 and Repo-GS>EcR RNAi #1 flies for three replicates, N=43-63, one-way ANOVA analysis with Turkey post-hoc test was used for (C), and Kruskal-Wallis test with Dunn’s multiple comparisons test was used for (D). (E-H) The average sleep bout number and average sleep bout length of the nSyb-GS>EcR RNAi #1 and Repo-GS>EcR RNAi #1 flies for all three replicates. Daytime sleep data are quantified in the left panels, and nighttime sleep is quantified in the right panels of each group. One-way ANOVA analysis with Turkey post-hoc test was used for (E-F) and Kruskal-Wallis test with Dunn’s multiple comparisons test was used for (G-H). (I-K) The rhythmicity, period, and relative FFT power analysis of nSyb-GS>EcR RNAi #1 and Repo-GS>EcR RNAi #1 flies and controls assayed for locomotor activity rhythms in constant darkness. Flies used for analysis in (J, K) are rhythmic flies from (I), and experimental files are compared with their GeneSwitch and RNAi control flies. Bar graphs show mean + SEM, and p values for each comparison were calculated using the Kruskal-Wallis test with Dunn’s multiple comparisons test. ns=not significant, p>0.05, *p<0.05, **p<0.01, ***p<0.001, ****p<0.0001.

Further sleep episode analysis showed that neuronal or glial-specific knockdown of EcR or E75 reduces sleep amount by fragmenting sleep, with larger effects seen with the glial-specific knockdown (**Figure2 and FigureS1**). Glial EcR and E75 knockdown flies had shorter average sleep bout lengths during the day and night and more sleep bouts at night, indicating that their sleep is fragmented (**Figure2 and FigureS1**). Comparison of RU486-treated flies with vehicle-treated controls confirmed that adult-specific knockdown of EcR is sufficient to reduce sleep in both neuronal and glial populations (**FigureS2**). We also tested the function of the recently identified ecdysone membrane transporter EcI, whose expression is necessary for ecdysone uptake by cells (Okamoto et al., 2018). As with EcR, glial knockdown of EcI reduced overall sleep levels and led to more fragmented sleep (**FigureS3E–S3H**). Knockdown of EcI in glia seemed to have a weaker effect on sleep than knockdown of EcR and E75, perhaps because EcI is one of the most highly expressed genes in glia and may be insufficiently knocked down by RNAi (DeSalvo et al., 2014). In general, all these data strongly indicate that ecdysone signaling acts in the brain, particularly in glia, to modulate sleep.

To assay flies for free-running circadian rhythms, we monitored them under constant darkness (DD) conditions. In DD, half of the nSyb-GS EcR knockdown flies and 90% of Repo-GS EcR knockdown flies became arrhythmic, but the remaining rhythmic flies showed no difference from controls in their period or FFT (Fast Fourier Transform) values that are a measure of rhythm strength (**Figure 2I–2K**). Our finding that EcR functions specifically in glia to promote circadian locomotor rhythms and sleep is novel and consistent with previous work showing that E75 or EcR knockdown in central clock cells causes some locomotor arrhythmicity, but more dramatic phenotypes result from knockdown with drivers such as *tim^27^-Gal4* that additionally express in glia (Kumar et al., 2014).

Next, we overexpressed EcR and found that, in contrast to our knockdown experiments, overexpression in neurons and glia had no effect on total sleep, but Repo-GS>EcR-c flies showed a consistent phase shift at dusk (**FigureS3A–3D**). Thus, effects of EcR on sleep and rhythms are largely through glia, but not neurons. Regarding the lack of sleep phenotypes from overexpression, it is possible that sleep-relevant ecdysone signaling is saturating in both neurons and glia and increasing EcR does not confer new sleep-promoting functions. However, since exogenous ecdysone feeding promotes sleep (Ishimoto and Kitamoto, 2010a), it is also possible that the EcR ligand ecdysone is rate-limiting when EcR is overexpressed.

**Figure3:**
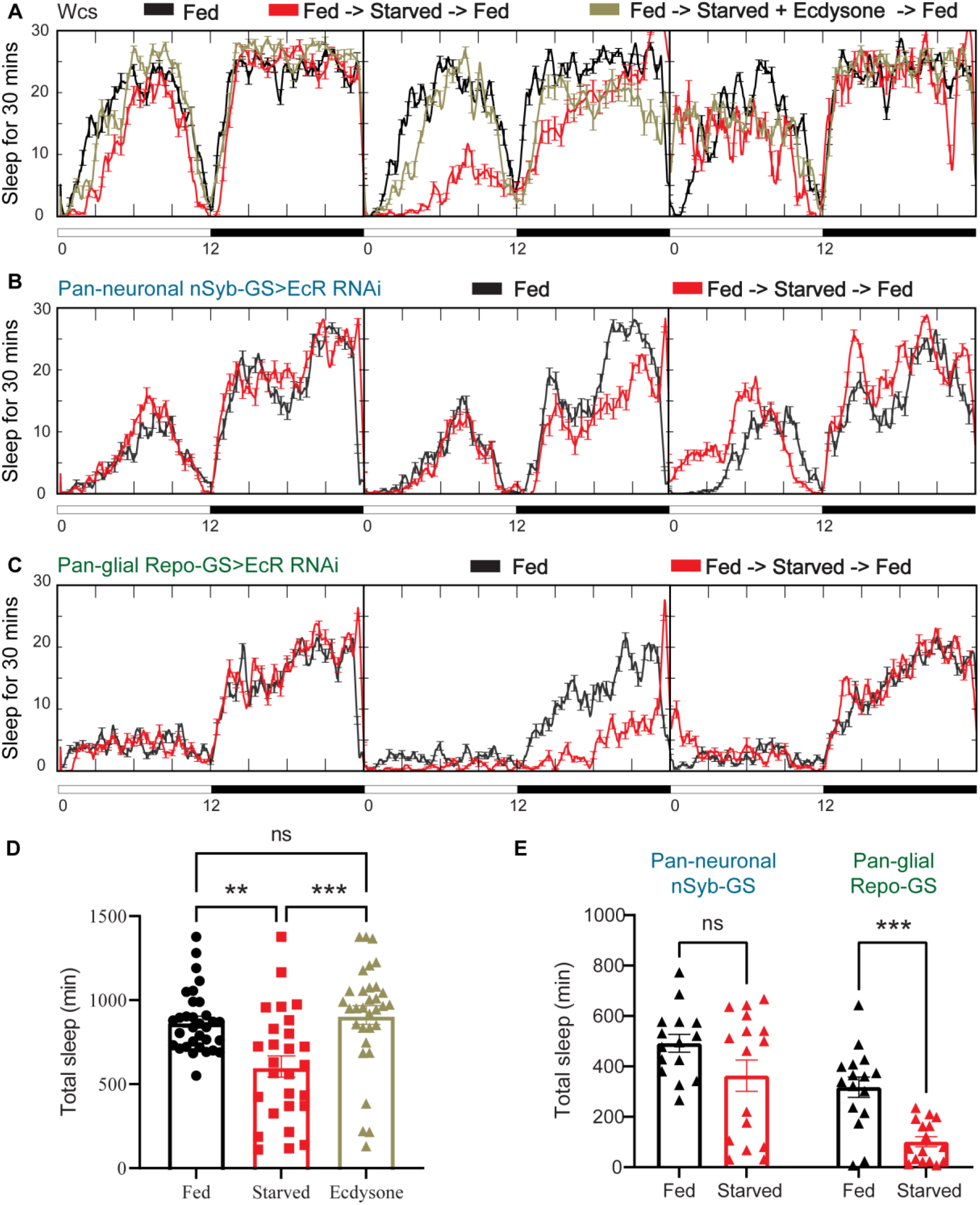
Ecdysone feeding prevents sleep loss in response to starvation, while EcR KD exacerbates starvation-induced sleep loss. Sleep was monitored for three days under each of the following three conditions--continuous feeding for three days or with an intervention on the second day, which consisted of either starvation or starvation accompanied with the feeding of 0.2mM ecdysone. The assay was conducted in (A) wild-type flies (*w*CS) and (B-C) nSyb-GS and Repo-GS EcR RNAi #1 flies. N=15-16 for each genotype, and the experiment was repeated multiple times with a one- or two-day starvation period and different ecdysone concentration. Quantification of sleep for the ZT0-ZT23h interval of the second day is shown for (D) *w*CS female flies and (E) nSyb-GS and Repo-GS EcR RNAi #1 female flies. ZT0-ZT23 sleep data are shown as flies were flipped back to locomotor tubes with food during the last hour (ZT23-24). Bar graphs show mean + SEM and ns=not significant, p>0.05, * p<0.05, ** p<0.01, *** p<0.001, **** p<0.0001. P values for comparisons between two groups were based on the Mann-Whitney test (E), and P values for comparison between three groups were calculated by one-way ANOVA with Turkey post-hoc test (D).

### Starved flies are particularly sensitive to loss of ecdysone signaling in glia

Sleep changes under stress conditions, and ecdysone is implicated in stress responses (Hill et al., 2018; Ishimoto and Kitamoto, 2011; Williams, 2019), raising the possibility that ecdysone mediates the effects of stressors on sleep. Starvation is an example of a stressor that raises ecdysone levels in female flies (Terashima et al., 2005). Although its effect on sleep goes in the opposite direction, i.e., starvation reduces sleep, we asked whether ecdysone is relevant for effects of starvation (Keene et al., 2010); for instance, does downregulation of ecdysone signaling in specific cells (despite overall increased ecdysone) account for decreased sleep during starvation? We first asked if starved flies respond to ecdysone by feeding them the exogenous bioactive form of ecdysone, 20E (Ishimoto and Kitamoto, 2010b). Exogenous ecdysone abrogated sleep loss caused by starvation, indicating that EcRs are functional in sleep-relevant cells under these conditions (**Figure 3A**). Consistent with this, we found that mRNA levels of ecdysone-responsive genes—EcR, EcI, E75, and E74— in the fly brain are not altered after one day of starvation (**FigureS5D**). Interestingly, starved flies fed 20E showed rebound sleep when they were returned to food, even though these flies did not lose sleep during starvation, indicating that 20E does not rescue the need for sleep that builds up during starvation (Regalado et al., 2017).

To determine whether increased sleep after 20E feeding is mediated by neurons or glia, we knocked down EcR in each of those two cell types. Sleep increased following 20E administration in nSyb-GS>EcR RNAi flies, but not Repo-GS>EcR RNAi flies. As the Repo-GS>Wcs flies also did not respond to 20E feeding (**Figure S4A**), we cannot draw conclusions about the necessity of ecdysone receptor expression in glia for effects of exogenous ecdysone. However, neuronal ecdysone signaling appears to be dispensable for this purpose.

Although exogenous ecdysone increased sleep in starved flies, it is still possible that endogenous neuronal/glial EcR signaling is disrupted in these flies, for example less ecdysone may enter the brain despite the overall elevated ecdysone level, in which case the knockdown of EcRs would have little effect. Surprisingly, we found that in starved flies while neuronal EcR knockdown did not significantly reduce sleep, EcR knockdown in glia very rapidly and profoundly reduced sleep (**Figure 3B–3E**). Thus, glial ecdysone signaling is particularly relevant for maintaining sleep during starvation. We also examined the response to a stressor that increases sleep, heat shock, and found that sleep induced by heat shock was not affected by EcR knockdown in neurons or glia (**Figure S4B**).

### Ecdysone cycles in the periphery and has higher action on the brain at night

Given that ecdysone modulates rhythmic behavior, the question arises whether it is under circadian regulation. We used multiple methods, including a genetic reporter (hs-Gal4-EcR.LBD), mass spectrometry, and an ecdysone ELISA (enzyme-linked immunosorbent assay) kit, to determine if 20E cycles in the brain over a day-night cycle (Ishimoto and Kitamoto, 2010b). However, through ELISA analysis, we found that the 20E level, which likely depends upon integrating many signals, is significantly higher at ZT14 than ZT2 in the fly periphery but not in the fly brain (**Figure S5A**). In the fly brain, even genetic reporters and mass spectrometry approaches failed to detect any significant differences in 20E levels. Our inability to detect a cycle in the brain, despite a previously reported peak at ZT12 in the fly head (Ishimoto and Kitamoto, 2010b), could reflect the rapid metabolism of 20E following its action on brain tissues. Alternatively, changes of 20E levels in the fly brain may be too low to be detected (Vafopoulou and Steel, 2012). In support of circadian regulation of ecdysone signaling, mRNA levels of ecdysone responsive E75 isoforms cycle with a peak at ZT12 in the fly head, suggesting that ecdysone acts rhythmically in the fly brain (**Figure S5B**). Given that E75 isoforms do not peak at ZT12 in the fly periphery, we suggest that elevated ecdysone at this time enters the brain instead of acting on peripheral tissues. To determine whether ecdysone enters the brain more efficiently at night, we examined whether E75 isoforms show differential responses to peripheral injection of 20E at ZT 6 versus ZT18. We normalized the change in E75 expression in the brain to the change in the body to account for the inconsistency of injection and found that the mRNA levels of both E75A and E75B increased more at ZT18, compared to ZT6, after 20E injection, 8.183 times more for E75A and 1.425 times more for E75B (**Figure S5C**). This result indicates that peripheral ecdysone has higher action in the brain at night.

### EcR functions in cortex glia to affect sleep

Given the relevance of glia to EcR effects on sleep, we focused on uncovering its function in glial cells. We first assayed the expression of EcR in adult brains. Consistent with previous reports, antibody staining in the adult fly brain showed that EcR is broadly expressed, but largely in the cortex glia. To confirm the localization to cortex glia, we used two separate cortex glia drivers, *GMR77A03-Gal4* and *Np2222-Gal4,* to drive the expression of mCD8-RFP. Co-staining these brains with an antibody against EcR confirmed that EcR is expressed primarily in the cortex glia (**Figure4A**). When EcR was knocked down in glia with Repo-GS, EcR antibody staining was greatly diminished. However, signals were preserved when EcR was knocked down in neurons, leading us to infer that glia, in particular cortex glia, are the major site of expression (**Figure4B**). Although not evident in these images, EcR expression was also detected in surface glia of the blood brain barrier (BBB) (data not shown).

**Figure 4:**
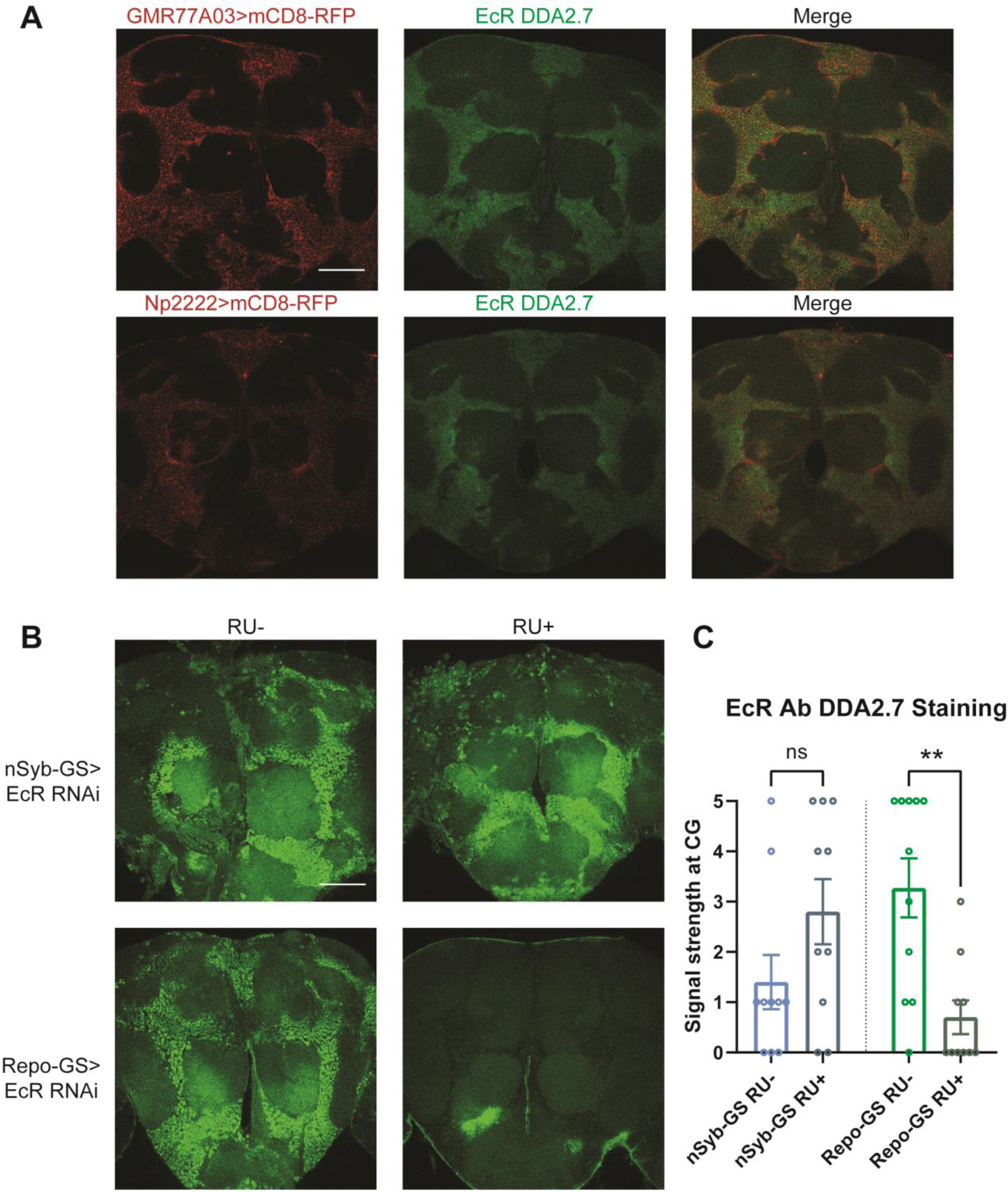
EcR is expressed in cortex glia. (A) EcR antibody DDA2.7 staining overlaps with reporter expression driven by cortex glia drivers (GMR77A03>mCD8-RFP and Np2222>mCD8-RFP), indicating that EcR is expressed in the cortex glia. EcR antibody DDA2.7 staining also shows that EcR can be nuclear or cytoplasmic. (B) EcR antibody DDA2.7 staining is preserved and still observed in the nSyb-GS>EcR RNAi flies but is almost eliminated in the Repo-GS>EcR RNAi flies compared with the vehicle control flies. N=10 per group. Scale bar: 100 μm. (C) Quantification of antibody staining in the fly brains’ cortex glia layer. Signal strength was scored manually based on the fluorescence intensity in the cortex glia region. Repo-GS>EcR RNAi #1 flies have significantly reduced staining based on an unpaired parametric Student t-test.

To determine whether sleep phenotypes correlate with the expression pattern of EcR, we then knocked down EcR constitutively in different glial subpopulations by crossing subglial Gal4 lines with the EcR RNAi #1 line that targets all EcR isoforms. However, EcR knockdown with most of these Gal4 lines caused eclosion failure, indicating the importance of EcR in glial cells during development. Three Gal4 lines produced viable adult flies: one cortex glia-specific driver *GMR77A03*, one astrocyte-like glia-specific driver Eaat-1, and one ensheathing glia-specific driver *GMR56F03.* Of these three Gal4 lines, sleep was only reduced when EcR was knocked down using the cortex glia-specific driver (**Figure5A, 6C**).

**Figure5:**
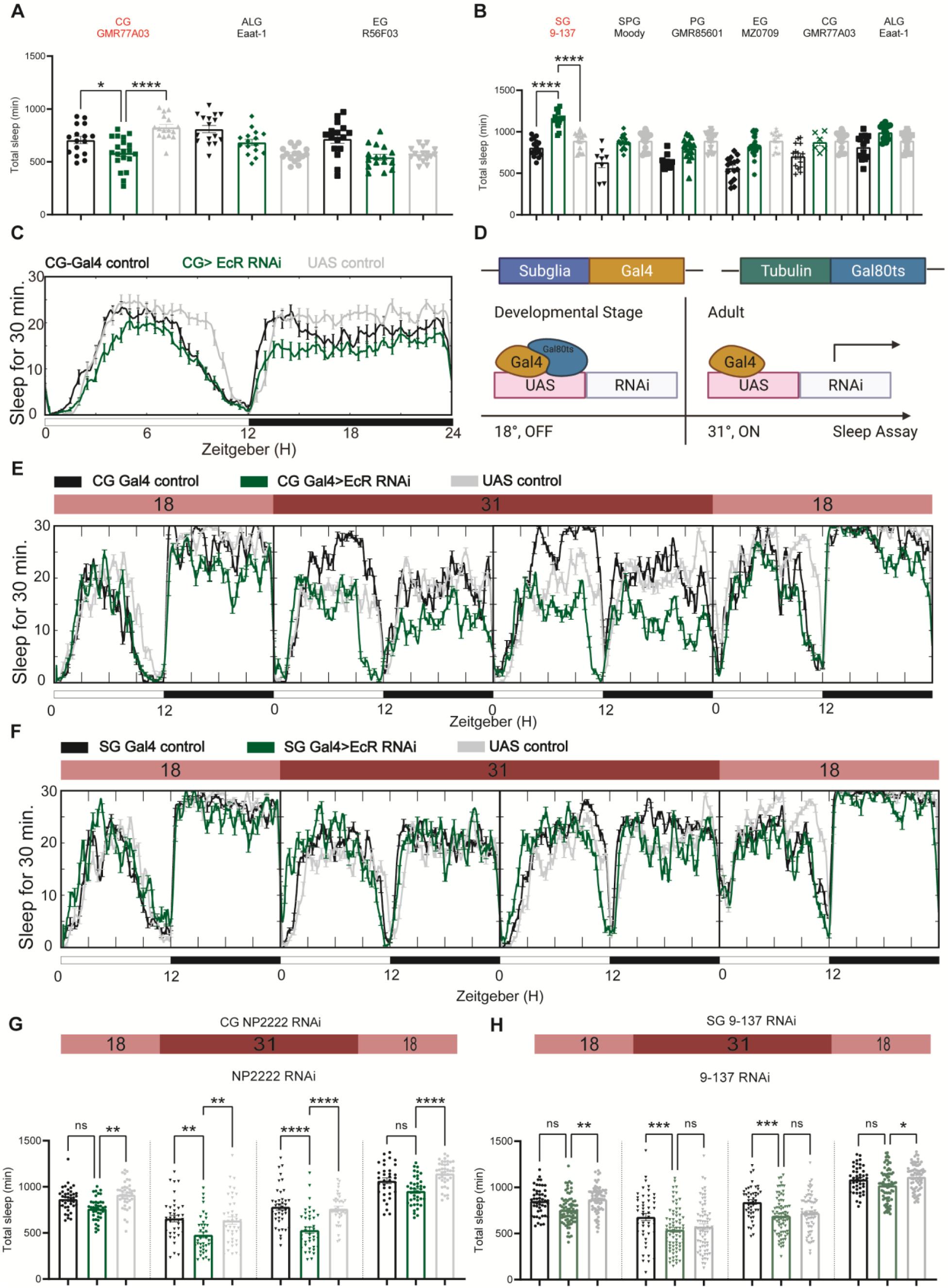
EcR functions in cortex glia to affect sleep. (A) Multiple constitutive Gal4 drivers labeling different sub-glial populations were crossed to EcR RNAi flies. Only GMR77A03 for cortex glia (CG), Eaat-1 for astrocyte-like glia (ALG), and GMR56F03 for ensheathing glia (EG) drivers produced viable adult progeny flies, and only GMR77A03>EcR RNAi #1 flies showed reduced total sleep compared with control flies. Green chart columns are experimental groups, and neighboring black and grey columns are Gal4 and UAS flies controls respectively. N=16-20 per genotype. (B) Overexpression of EcR by sub-glial Gal4 drivers—9-137-Gal4 to drive expression in the surface glia (SG), Moody-Gal4 for subperineurial glia (SPG), GMR85G01 for perineurial glia (PG), MZ008-Gal4 for ensheathing glia (EG), GMR77A03 for cortex glia (CG), and Eaat-1 for astrocyte-like glia (ALG). Only overexpression of EcR in the surface glia promotes sleep. N= 16-24 per genotype. (C) Representative sleep traces of the cortex glia GMR77A03>EcR RNAi #1 flies. (D) Gal4/Tubulin-gal80ts was used to achieve adult-specific knockdown of EcR in different sub glial populations. Under permissive temperature, Gal80ts inhibits Gal4 activation of UAS, but under restrictive temperature, Gal80ts is inactivated, and genes under the regulation of UAS are expressed. (E-F) Sleep traces resulting from EcR knockdown in the cortex glia using NP2222-Gal4/tubulinGal80ts and surface glia using 9-137-Gal4/tubulinGal80ts. F1 progeny flies were kept at 18 degrees for one day, and then the temperature was switched to 31 degrees to inactivate the Gal80ts and thus achieve knockdown of EcR over the following two days. Subsequently, temperatures were decreased back to 18 degrees. (G, H) show quantification of total sleep of all EcR knockdown flies in the cortex glia using NP2222-Gal4/tubulinGal80ts and surface glia using 9-137-Gal4/tubulinGal80ts, N =34-77 per genotype. Total sleep of each genotype was calculated and compared to controls for the above four days. Bar graphs show mean + SEM, ns=not significant, p>0.05, * p<0.05, ** p<0.01, *** p<0.001, **** p<0.0001. P values for each comparison were calculated by one-way ANOVA analysis with Tukey post-hoc test.

**Figure6.**
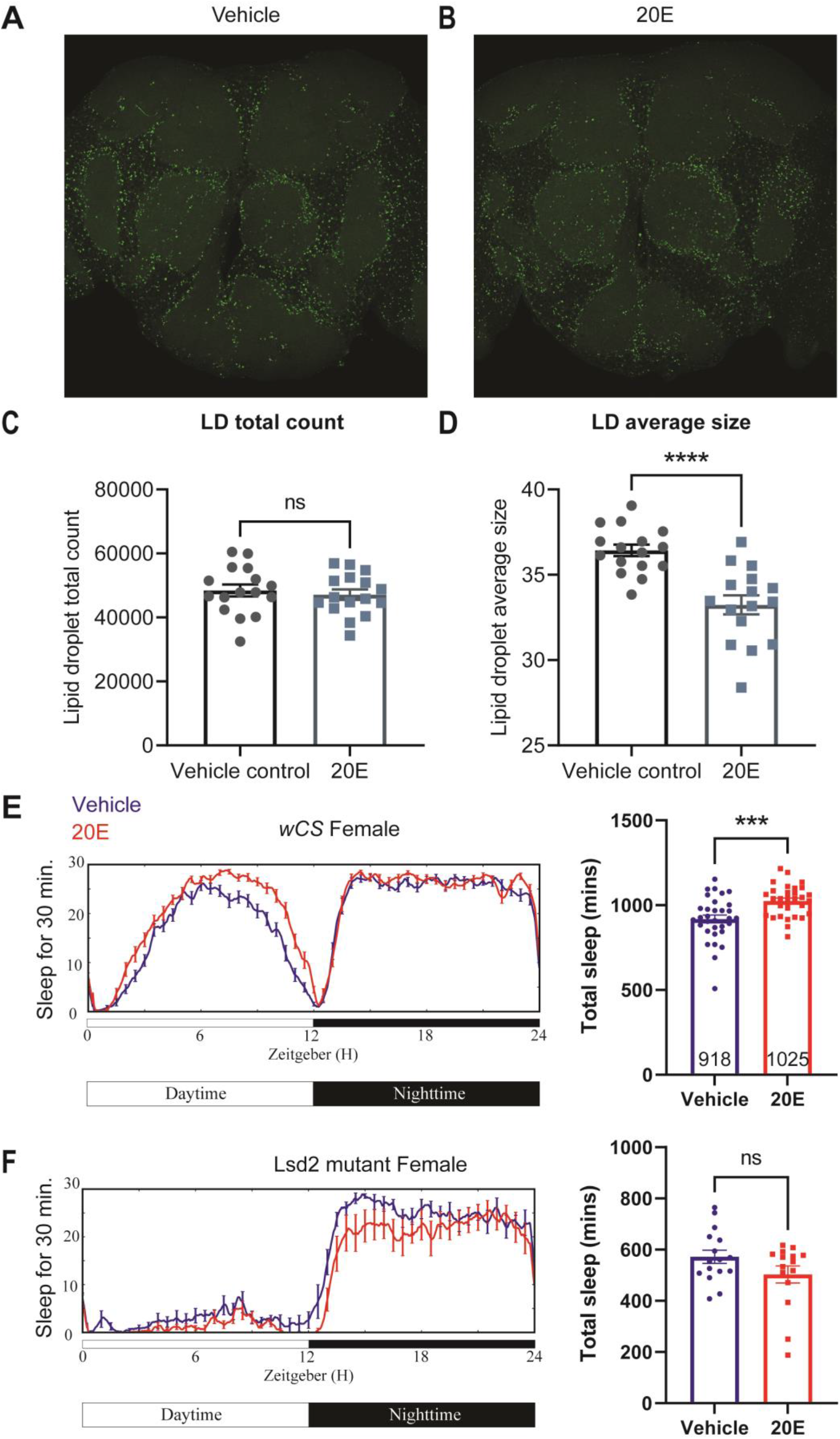
Lipid metabolism mediates the effects of 20E on sleep. (A-B) Lipid droplet (LDs) staining of representative brains from flies treated with vehicle or 0.5mM 20E. A z-stack slice that shows the maximal structure of the cortex glia was selected, and brightness and contrast were auto-adjusted by ImageJ for better visualization. LDs are stained by the lipophilic dye BODIPY 493. (C-D) 0.5 mM 20E treatment does not significantly affect the total count of LDs but likely mobilizes lipids to lead to smaller LDs. LD count and size were analyzed and calculated using ImageJ as detailed in the method session. N= 16 per group from two independent repeats. Statistical comparisons used unpaired parametric Student t-test. Bar graphs show mean + SEM and ns=not significant, p>0.05, * p<0.05, ** p<0.01, *** p<0.001, **** p<0.0001. (E-F) A representative sleep trace of *w*CS flies with vehicle control or 0.2mM ecdysone. As previously reported, total sleep increases with 20E treatment, N =32 per group. (B) The sleep-promoting effect of the 20E is lost in the *lsd2* mutant flies, N =16 per group from two separate experiments.

To avoid issues of lethality and to distinguish between developmental and adult-specific roles, we used the temporal and regional gene expression targeting (TARGET) system to repress Gal4 transcription in the larval stage, after which the flies were transferred to a (restrictive) elevated temperature to degrade temperature-sensitive GAL80^ts^ and allow expression of Gal4 (**Figure 5D**) (McGuire et al., 2004). At the (permissive) low temperature, when TUB-GAL80ts blocks Gal4 expression, *NP2222*-Gal4>EcR RNAi flies trended towards reduced sleep but sleep significantly decreased at the restrictive temperature when RNAi expression was activated to knock down EcR in cortex glia (**Figure 5E and 5G**). Knockdown of EcR with the 9-137 Gal4 driver, which targets all glia of the BBB, did not affect sleep (**Figure 5F and 5H**), and nor did knockdown of EcR in ensheathing or astrocyte glia (**Figure S6**), suggesting that effects of EcR on sleep are restricted to specific glia, with cortex glia being the primary physiological site of action.

We also overexpressed EcR using the same set of sub-glia drivers and found that only flies expressing EcR with the 9-137-Gal4 driver had elevated sleep (**Figure 5B**). These flies were also resistant to sleep deprivation, indicating enhanced sleep need. Restricting overexpression to the adult stage with TubGal80 did not affect sleep, suggesting that the sleep-promoting effect observed in 9-137-Gal4 flies is developmental. (**Figure S6D**).

### Ecdysone modulates sleep by mobilizing lipid droplets in glial cells

The cortex glia, especially the superficial cortex glial cells, are enriched in lipid droplets in the third instar larval stage, and ecdysone signaling can mobilize these lipids when needed (Kis et al., 2015). To determine whether ecdysone in adult flies has a similar effect on lipids, we assayed lipid droplets (LDs) in glial cells of flies fed ecdysone. We measured lipid droplets (LDs) in the fly brain by BODIPY staining and found that LDs, which accumulate mostly in the cortex and ensheathing glia, were significantly smaller in flies fed ecdysone, relative to control groups, at Zeitgeber Time (ZT) 12. At the same time, the total count was not affected (**Figure 6A–6D**). We next asked if this effect on lipids was relevant for sleep induction by ecdysone by measuring the ecdysone response of a lipid storage mutant (Kamoshida et al., 2012). The Lipid storage droplet 2 (LSD2) protein modulates lipid accumulation and response to starvation, so *lsd2* mutants are lean and sensitive to starvation (Thimgan et al., 2010). We found that ecdysone does not promote sleep in *lsd2* mutants (**Figure 6E–6F**), These results suggest that ecdysone affects sleep by modulating LDs in glial cells, especially cortex glia.

## Discussion

While sleep traditionally has been regarded as a neuronally-driven behavior, glial cells can regulate sleep by affecting neuronal activity and possibly even mediate functions attributed to sleep, such as waste clearance, nutrient transfer, and repair (Cai et al., 2021; Donlea et al., 2014; Keene et al., 2010; Ortega, 2016; Seidner et al., 2015; Singh and Donlea, 2020; Stanhope et al., 2020; Yildirim et al., 2019). We show here that glia are also a major target of steroid hormone signaling to regulate sleep. In addition, our data support an important role for lipid metabolism in controlling sleep.

In *Drosophila,* ecdysone is the major steroid hormone, and it plays an essential role in regulating metamorphosis and molting (Yamanaka et al., 2013a). Ecdysone also affects the development of the nervous system and neuronal remodeling, processes that may involve glial function (Yu and Schuldiner, 2014). During development, ecdysone is synthesized from dietary cholesterol in the prothoracic gland (PG) and in other peripheral tissues that control molting and eclosion in larvae and pupae, but it declines during pupa-adult transitions and remains low in adults (Yamanaka et al., 2013b (Kannangara et al., 2021)). Nevertheless, it is implicated in adult functions such as memory formation and stress resistance (Ishimoto and Kitamoto, 2011). Ecdysone was also shown to affect sleep (Ishimoto and Kitamoto, 2011), raising questions of the mechanism by which it does so and the cells on which it acts in adults. Our finding that ecdysone regulates sleep through metabolic mechanisms in glia is consistent with other metabolic functions attributed to it; for instance, it controls developmental transitions in response to nutrient signals (Christensen et al., 2020; Lee et al., 2000; Uyehara and Mckay, 2019; Xu et al., 2020; Yamanaka et al., 2013b). Indeed, ecdysone may affect the whole-body lipid profile to modulate physiology and behavior (Sieber and Spradling, 2015).

The gonads, in particular the ovaries in females, are thought to be the major source of the ecdysone in adults (Ahmed et al., 2020). However, several other tissues also express ecdysone biosynthesis genes (Li et al., 2022), so multiple peripheral tissues could contribute to circulating ecdysone levels. Consistent with this notion, our efforts to observe sleep changes by disrupting the synthesis of ecdysone in any one of the following peripheral tissues— gut, fat body and ovary— failed (Data not shown). We speculate that knockdown of biosynthetic enzymes in any one peripheral tissue does not have a significant effect because ecdysone can still be derived from other tissues. Notably, knockdown in the brain did not have any effect either. Also, Halloween genes have very little to no expression in the fly brain based on fly brain single cell sequencing (Davie et al., 2018), suggesting that ecdysone or 20E derives from the periphery to modulate sleep via glia.

In *Drosophila,* around 10%-15% of cells in the brain are glial cells belonging to one of five distinct groups: cortex glia, astrocyte-like glia, ensheathing glia, perineurial, and subperineural glia (Edwards and Meinertzhagen, 2010; Yildirim et al., 2019). Perineurial and subperineural glia together serve as the blood-brain barrier regulating permeability, controlled by the circadian system (Zhang et al., 2018). BBB glia also have a role in sleep, such that endocytosis in these cells increases during sleep and depends upon the prior duration of wakefulness (Artiushin et al., 2018). Astrocyte-like glia have a distinctive shape that allows remarkably close physical contact with synapses and is thought to be important for the clearance of neurotransmitters in the synaptic space (Freeman, 2015). In support of this, arylalkylamine N-acetyltransferase 1 (AANAT1), which acetylates and inactivates monoamines, acts in astrocytes to affect sleep (Davla et al., 2020). Also, calcium signaling in astrocytes appears to contribute to sleep need (Blum et al., 2021). Cortex glia cells encapsulate neuronal cell bodies and provide nutrients to neurons (Doherty et al., 2009). Cortex and ensheathing glia accumulate most lipid droplets in the brain, and fatty acid-binding protein (Fabp), which promotes sleep, is expressed in both glial populations (Gerstner et al., 2011). Thus, cortex and ensheathing glia are important for metabolic homeostasis of the fly brain, including nutrient transfer and balance between neurons and glia (Bittern et al., 2021). In addition, the effects of Amyloid precursor protein (App), which regulates the production and deposition of toxic amyloid peptides, on sleep are mediated by cortex glia (Luna et al., 2017).

We show here that the ecdysone signaling pathway functions in adult cortex glia and neurons to affect circadian locomotor rhythms and sleep. The specific cellular targets through which ecdysone regulates sleep were previously not known. Additionally, while ecdysone can regulate circadian rhythms through neuronally-expressed receptors (Kumar et al., 2014), we find that knockdown of glial ecdysone receptors results in more severe behavioral arrhythmicity than neuronal knockdown (**Figure 2I-K**). As *Drosophila* astrocytes regulate circadian locomotor rhythms (Ng et al., 2011), they are attractive candidates for mediating the effects of ecdysone on rhythms. Our finding that EcR in specific sub-glial cells is also vital to the development of flies is surprising even though NHRs have well-documented roles in development. Ecdysone treatment upregulates the gene *glial cell missing (gcm),* while knockdown of EcR reduces the expression of *gcm in vitro,* suggesting that ecdysone influences the development and morphology of glial cells by regulating *gcm,* which determines the fate of the lateral glial cells (Wang et al., 2014). Thus, glia are an important target for biological actions of ecdysone on development and adult behavior.

We show that exogenous ecdysone mobilizes lipids accumulated in cortex glia to promote sleep. Since cortex glia cells surround and compartmentalize neuronal cell bodies, they may modulate neuronal activity by facilitating metabolite transfer to neurons. Aside from cortex glia, the only glial type to yield a phenotype with manipulations of ecdysone signaling is the surface glia. The sleep phenotype by EcR overexpression in surface glia may be non-physiological as a strong Gal4 line produces it, but it nevertheless suggests that the surface glia can support ecdysone signaling.

Our finding that lipid metabolism is important for ecdysone-induced sleep fits with increasing evidence of interactions between sleep and lipids. Loss of sleep alters the lipid profile across species, including in human peripheral blood (Davies et al., 2014; Hinard et al., 2012; Weljie et al., 2015). Conversely, lipids have been shown to regulate sleep; for instance, the *lsd2* mutant we used here, as well as a mutant lacking a lipase, affect rebound after sleep deprivation in *Drosophila* (Thimgan et al., 2010). Interestingly, specific lipids are also implicated as secreted sleep inducers (somnogens) that promote sleep following deprivation in mammals (Cravatt et al., 1995). Studies in worm also showed that sleep is associated with fat mobilization, and deficits in energy mobilization in sensory neuroendocrine cells cause sleep defects (Grubbs et al., 2020).

How ecdysone synthesis is regulated and how its function is integrated with innate and environmental changes needs further study. Juvenile hormone (JH), which works together with ecdysone during developmental stages, differentially affects sleep in male and female flies (Leinwand and Scott, 2021; Schwedes and Carney, 2012; Wu et al., 2018); it is reasonable to hypothesize that JH interacts with ecdysone in the context of sleep. The human ortholog of E75 is Reverb, which has broad effects on metabolism in peripheral tissues based on work in mice and humans, so the functions of EcR/E75 in peripheral tissues may also be linked to metabolism (Ding et al., 2021; Zhang, 2017). In addition, Reverb has high expression in mouse brain glial cells, where it could function to affect mouse sleep (Chi-Castañeda and Ortega, 2017).

In summary, the endocrine system and the circuitry underlying circadian rhythms and sleep are intertwined during developmental stages (AlDabal and BaHammam, 2011; Gamble et al., 2014; Morgan and Tsai, 2015), and we now find that ecdysone acts through specific glial cells to affect circadian rhythms and sleep in adults. Similar effects of the ecdysone downstream target E75 and the ecdysone importer implicate canonical nuclear hormone signaling in sleep regulation. The mechanism involves lipid droplet mobilization, emphasizing the importance of glia and lipid metabolism in sleep regulation. Our findings are likely just the tip of the iceberg concerning endocrine regulation of sleep. We expect this to be a rich area of investigation in the future, as peripheral effects on brain function are increasingly recognized.

## Acknowledgments

This work was supported by Howard Hughes Medical Institute (HHMI). We thank Anna Kolesnik, Joy Shon, Xanthe Heifetz Ament, and Kiet Luu for continuing technical assistance and all other members of the Sehgal lab for feedback, comments, and reagents. We thank Dr. Michael Welts and Dr. Naoki Yamanaka for generously sharing fly lines with us.

## Author contributions

Conceptualization, Y.J. Li and A.S.; Methodology, Y.J. L., P.H., and Z.F.Y.; Investigation, Y.J. L., P. H., and S.Z. Writing – Original Draft Y.J.L. and A.S.; Writing – Review & Editing, P.H., S.Z., and A.S.; Funding Acquisition, A.S.; Resources, Y.J.L., and A.S; Supervision, S.Z., and A.S.

---

**Figure S1:**
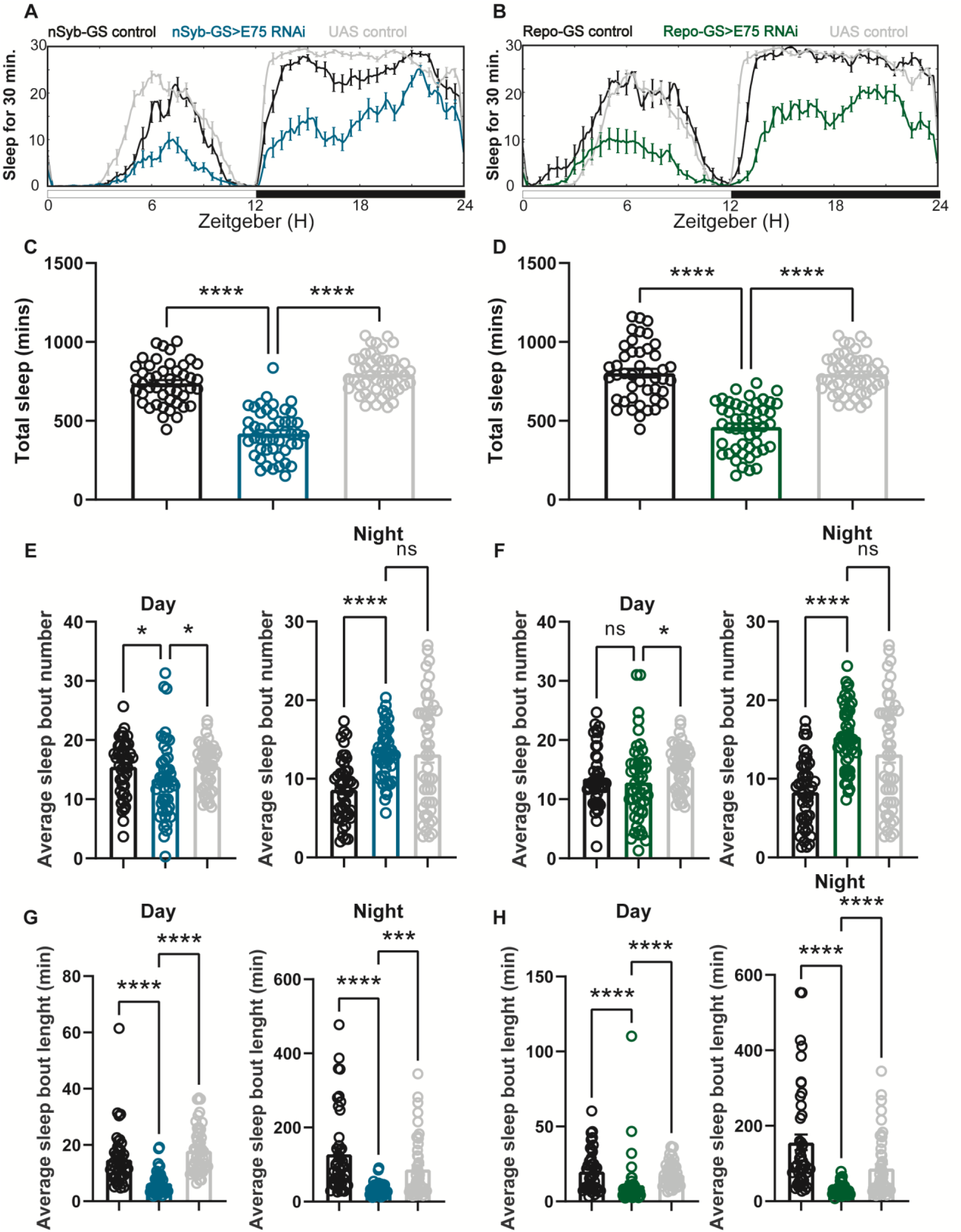
Baseline sleep phenotypes resulting from pan-neuronal and pan-glial knockdown of E75, the direct downstream target of EcR. (A-B) show representative sleep traces of nSyb-GS>E75 RNAi #1 and Repo-GS>E75 RNAi #1. N=12-16 per genotype per experiment. Experiments were independently repeated at least two times, and one experiment is shown here. (C-D) Total sleep of the nSyb-GS>EcR RNAi #1 and Repo-GS>EcR RNAi #1 flies shown in (A-B, p values for each comparison were calculated by one-way ANOVA analysis with the Tukey post-hoc test. (E-H) The average sleep bout number and average sleep bout length of the nSyb-GS>E75 RNAi #1 and Repo-GS>E75 RNAi #1 flies. Daytime sleep data are quantified in the left panels, and nighttime sleep is quantified in the right panels of each group. P values for each comparison in (E-H) were calculated by Kruskal-Wallis test with Dunn’s multiple comparisons test. Bar graphs show mean + SEM. ns=not significant p>0.05, *p<0.05, **p<0.01, ***p<0.001, ****p<0.0001.

**Figure S2:**
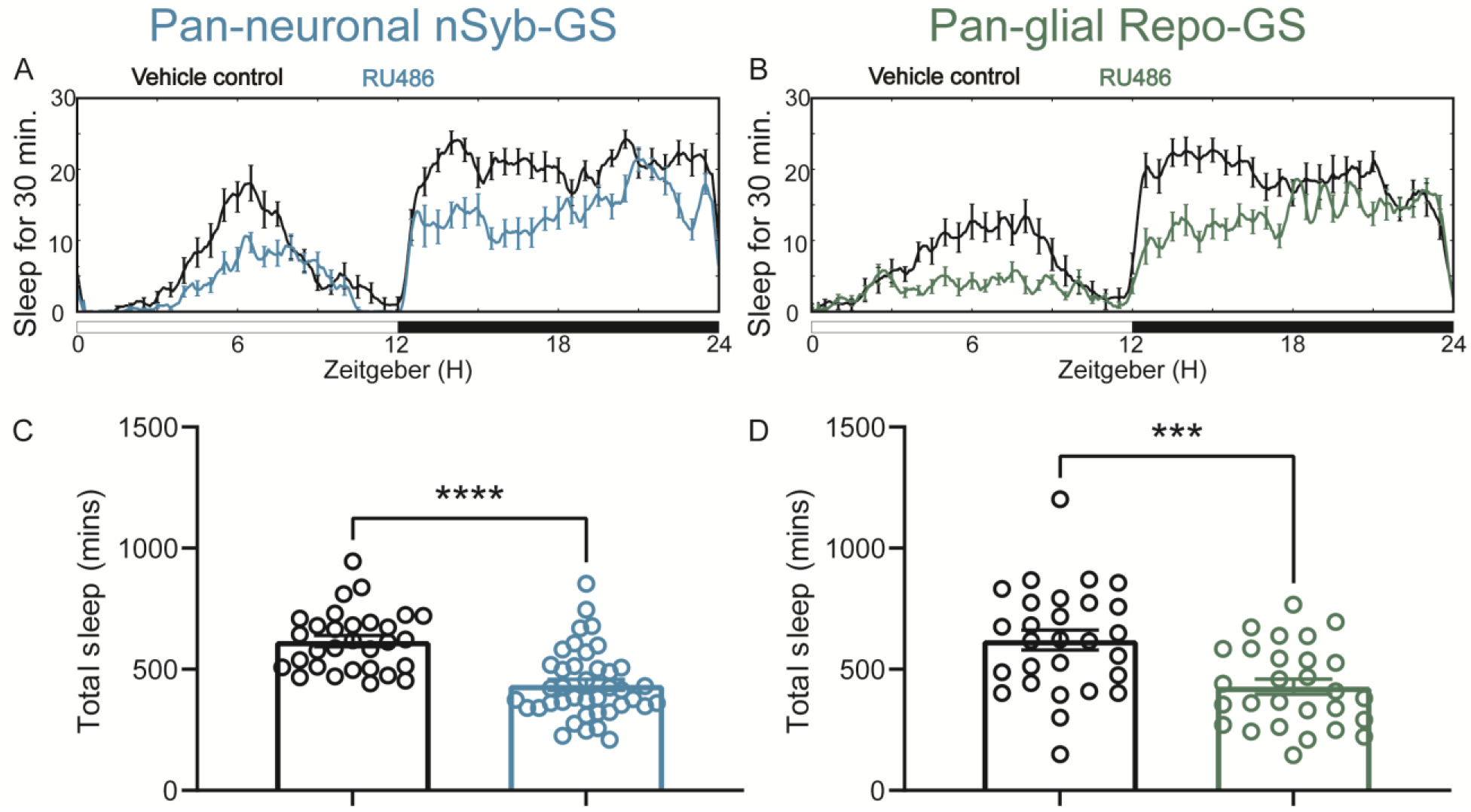
Adult-specific neuronal and glial knockdown of EcR reduces sleep. (A-B) show representative sleep traces of nSyb-GS>EcR RNAi #1 and Repo-GS>EcR RNAi #1 female flies with vehicle control or RU486; experiments were independently repeated at least two times. (C-D) Total sleep of the nSyb-GS>EcR RNAi #1 RNAi and Repo-GS>EcR RNAi #1 female flies with vehicle control or RU486, N=28-40 per genotype. Bar graphs show mean + SEM, and ns=not significant p>0.05, * p<0.05, ** p<0.01, *** p<0.001, **** p<0.0001. The comparison was calculated with an unpaired parametric Student t-test for Repo-GS groups and Mann-Whitney test for nSyb-GS groups.

**Figure S3:**
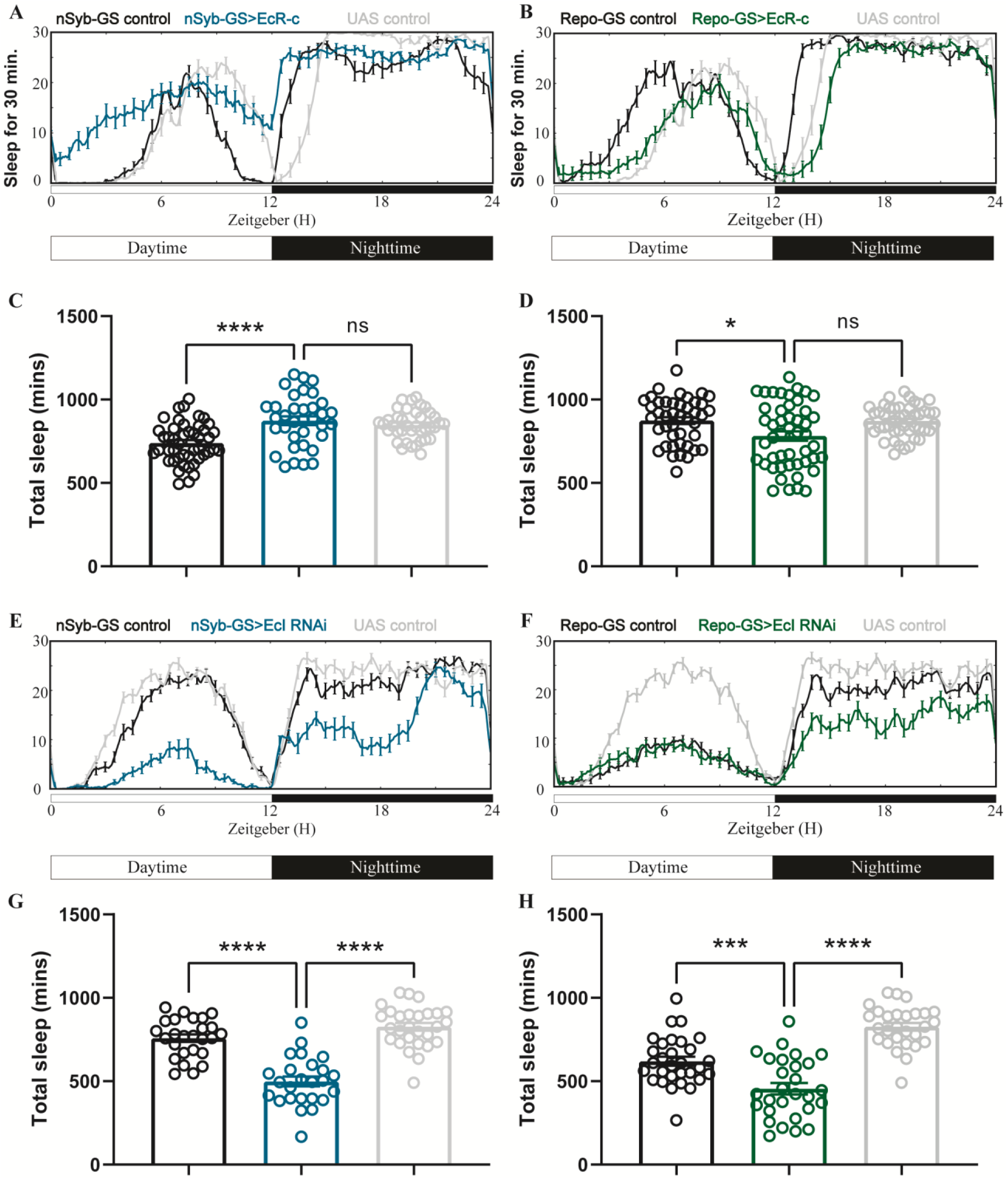
Baseline sleep phenotypes resulting from pan-neuronal and pan-glial overexpression of EcR common isoforms and knockdown of EcI, the membrane importer of ecdysone. (A-B) show representative sleep traces of nSyb-GS>EcR_c and Repo-GS>EcR_c flies, N=35-48 per genotype; data are based on three independent experiments. (C-D) Total sleep of the nSyb-GS>EcR_c and Repo-GS>EcR_c flies. (E-F) show representative sleep traces of nSyb-GS>EcI RNAi #1 and Repo-GS>EcI RNAi #1 flies, N=25-30; data are based on two independent experiments. (G-H) Total sleep of the nSyb-GS>EcI RNAi #1 and Repo-GS>EcI RNAi #1 flies. Bar graphs show mean + SEM; ns=not significant p>0.05, * p<0.05, ** p<0.01, *** p<0.001, **** p<0.0001. P values for each comparison were calculated by one-way ANOVA analysis with Turkey post-hoc test.

**Figure S4:**
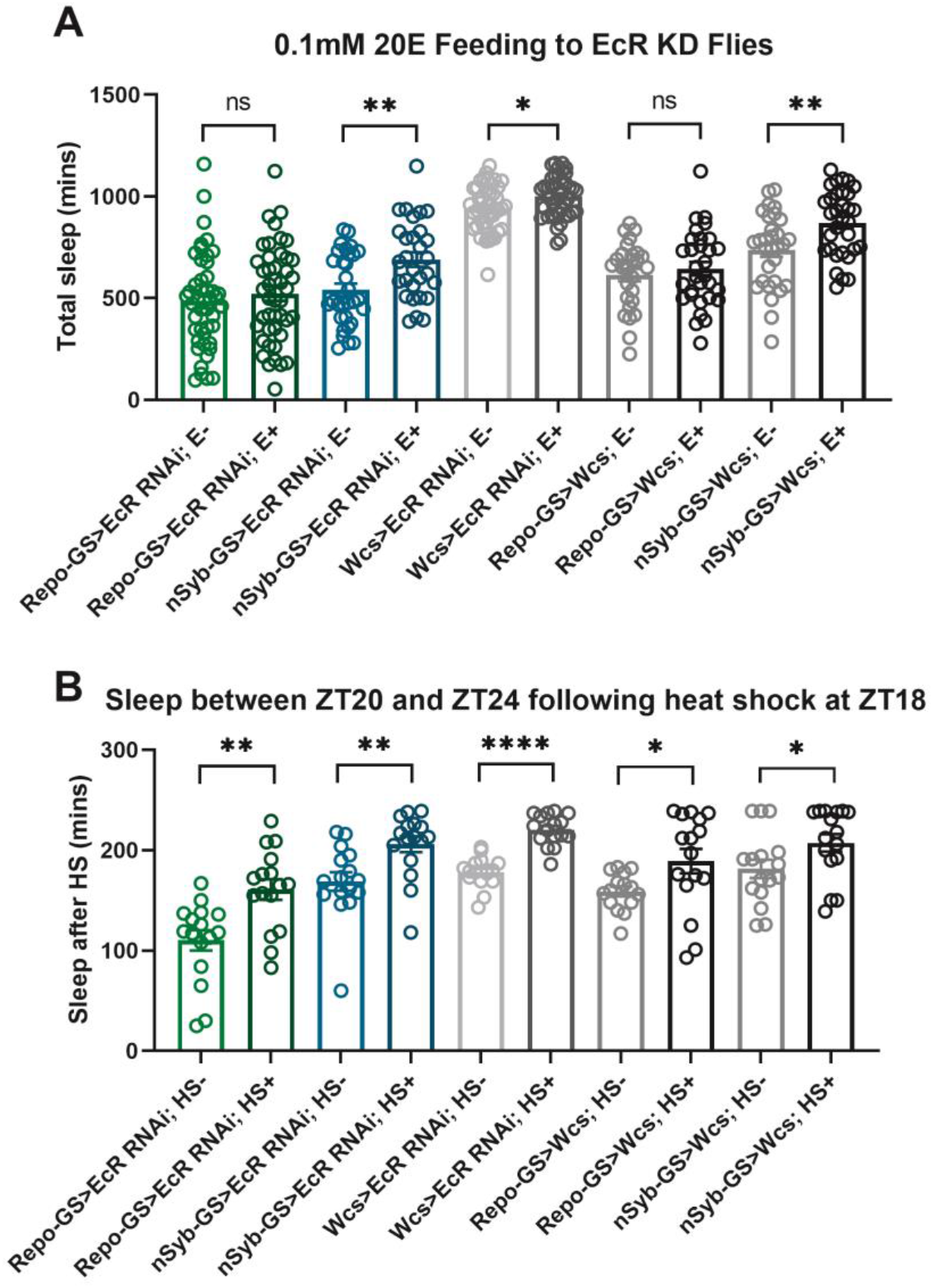
EcR disruption in neurons or glia does not affect the response to heat shock. (A) Total sleep of EcR RNAi #1 knockdown flies in both neurons and glia and their genetic control flies after 0.1mM 20E feeding, N=29-46, and experiments were repeated two or three times independently. (B) The last 4 hours’ sleep between ZT20-24 after heat shock for an hour at ZT18 of EcR RNAi # 1 knockdown flies in neurons or glia and their genetic control flies, N=16. Bar graphs show mean + SEM; ns=not significant p>0.05, * p<0.05, ** p<0.01, *** p<0.001, **** p<0.0001. P values for each comparison were calculated by one-way ANOVA analysis with Turkey post-hoc test.

**Figure S5:**
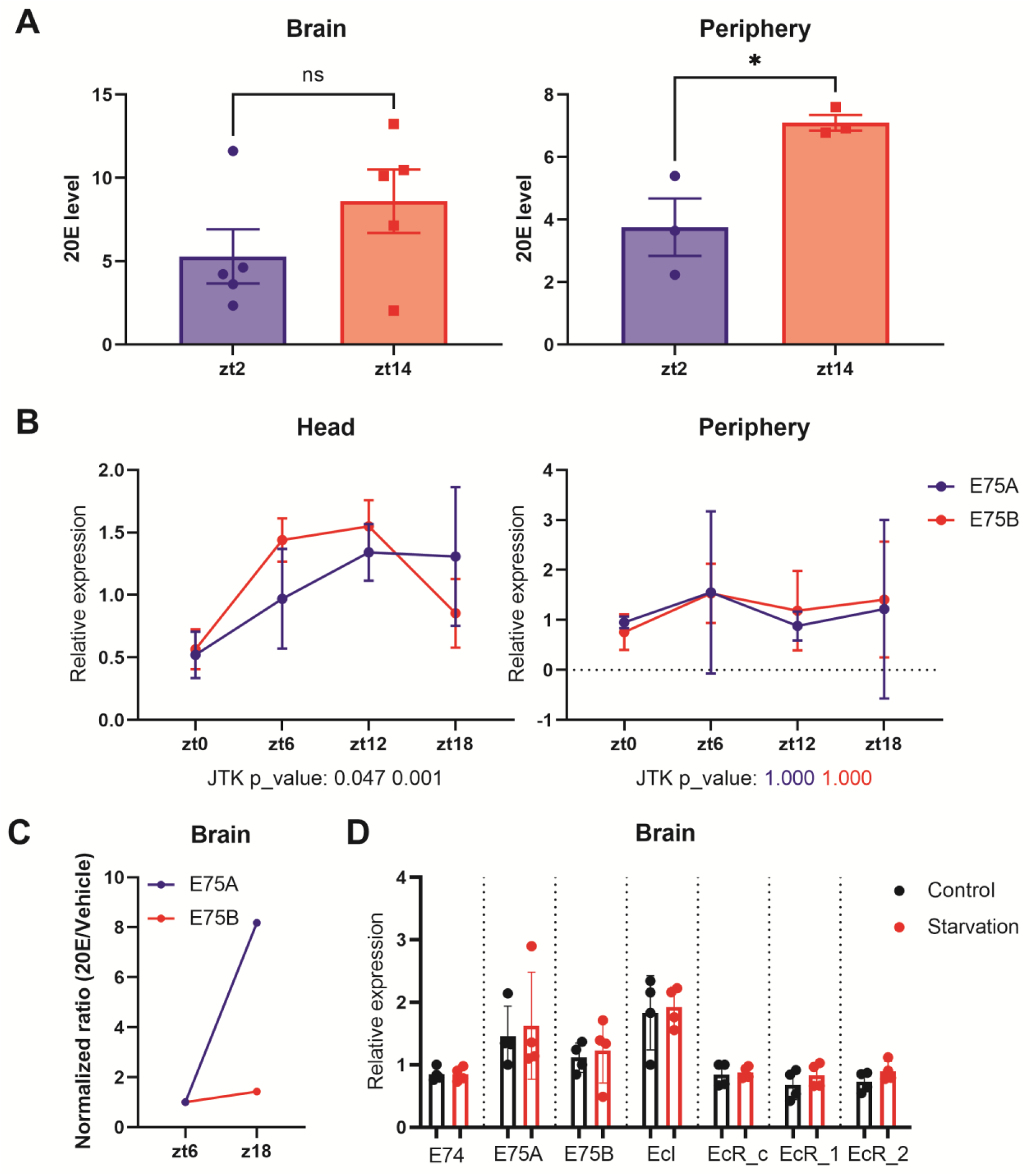
Ecdysone levels are higher in the fly periphery at ZT14 compared to ZT2. (A) Ecdysone level was measured in the fly’s whole body or brain using an ELISA kit. 20E levels are higher at ZT14 than ZT2 in the periphery. (B) A and B isoforms of the direct downstream target gene of ecdysone, E75, were measured by qPCR in fly heads or fly bodies at four different time points. Data indicate a significant rhythm with a mid-afternoon-evening peak in heads (JTK values, which are measures of cycling, are below the x-axis). (C) E75 isoform A and B mRNA levels in the fly brain following 20E/Vehicle injection at ZT6 or ZT18. The mRNA levels were measured 1 hour after flies were injected with 20E or vehicle control and changes seen with 20E (relative to control) in the brain at different time points were normalized to the changes in the body to control for inconsistency of the injection. (D) mRNA levels of EcR, EcI, E75, and E74 in the fly brains do not respond to 1 day of starvation. All comparisons’ p>0.05 in (C); thus, these are not shown. Bar graphs show mean + SEM, and ns=not significant p>0.05, * p<0.05, ** p<0.01, *** p<0.001, **** p<0.0001. P values for each comparison are calculated by unpaired parametric student t-test.

**Figure S6:**
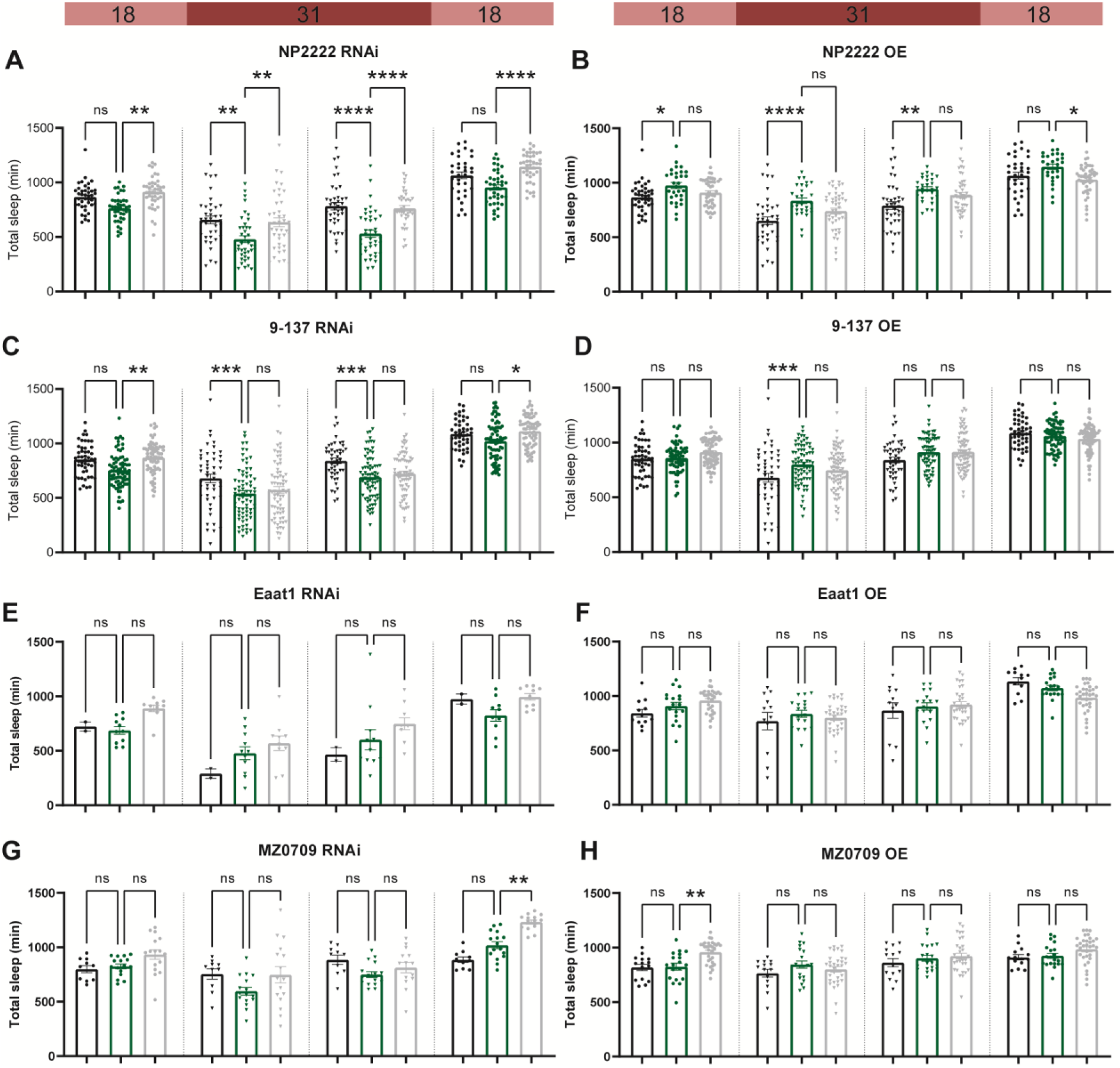
Adult-specific disruption of EcR in most sub glial populations does not affect sleep. Temperature-sensitive tubulin-Gal80ts were used to restrict the expression of Gal4 to adults. Sleep was monitored for one day at the permissive temperature of 18 degrees, then two days at the restrictive temperature of 31 degrees data, and then for another day at 18 degrees. EcR knockdown (RNAi) and overexpression (OE) were driven by: (A-B) A cortex glia driver (NP2222>EcR RNAi #1 and NP2222>EcR OE); (C-D) A surface glia driver (9-137>EcR RNAi #1 and 9-137>EcR OE). (E-F) An astrocyte-like glia driver (Eaat-1>EcR RNAi #1 and Eaat-1>EcR OE). (G-H) An ensheathing glia driver (MZ0709>EcR RNAi #1 and MZ0709>EcR OE). Bar graphs show mean + SEM; ns=not significant p>0.05, * p<0.05, ** p<0.01, *** p<0.001, **** p<0.0001. p values for each comparison were calculated by one-way ANOVA analysis with the Tukey post-hoc test.

**Table S1.** Pan-neuronal screening of NHRs in adult *Drosophila*

| Genotype | RNAi line | N | sleep TD<br>(E-Gal4) | sleep RD<br>(E-Gal4) | sleep RN<br>(E-Gal4) | sleep TD<br>(E-UAS) | sleep RD<br>(E-UAS) | sleep RN<br>(E-UAS) |
| --- | --- | --- | --- | --- | --- | --- | --- | --- |
| nSyb-GS>E75 RNAi GD | **E75 #1** | 47 | -309.4311044 | -119.49582 | -189.935 | -356.474 | -158.59 | -197.884 |
| nSyb-GS>Svp RNAi 44394 | Svp #1 | 16 | -259.140625 | -215.625 | -43.5156 | -227.203 | -183.109 | -44.0938 |
| nSyb-GS>Usp RNAi 36729 | **Usp #2** | 14 | -242.6629464 | -175.7567 | -66.9063 | -207.257 | -200.694 | -6.5625 |
| nSyb-GS>Ftz-f1 RNAi 33625 | **Ftz-f1 #2** | 15 | -240.3927083 | -135.61979 | -104.773 | -181.721 | -119.698 | -62.0229 |
| nSyb-GS>Hr3 RNAi 44399 | **Hr3 #1** | 15 | -220.8347222 | -107.76667 | -113.068 | -231.043 | -120.163 | -110.881 |
| nSyb-GS>Eip78c RNAi 44390 | **E78 #1** | 6 | -213.109375 | -168.24479 | -44.8646 | -180.453 | -134.182 | -46.2708 |
| nSyb-GS>E75 RNAi JF02257 | **E75 #2** | 32 | -165.7529762 | -83.93006 | -81.8229 | -125.188 | -128.312 | 3.125 |
| nSyb-GS>EcR RNAi 37059 | **EcR #2** | 8 | -157.047619 | -11.630952 | -145.417 | -25.2083 | -61.5208 | 36.3125 |
| nSyb-GS>Hr39 RNAi 27086 | **Hr39 #1** | 47 | -139.8584093 | -77.39653 | -62.4619 | -172.119 | -114.185 | -57.934 |
| nSyb-GS>Hnf4 RNAi 44398 | Hnf4 #1 | 21 | -133.9322917 | -88.30754 | -45.6248 | -218.912 | -162.531 | -56.3804 |
| nSyb-GS>Dsf RNAi 41707 | Dsf #2 | 16 | -130.78125 | -126.34375 | -4.4375 | -370.789 | -276.357 | -94.4313 |
| nSyb-GS>Dsf RNAi 29373 | Dsf #1 | 32 | -115.469122 | -29.573289 | -85.8958 | -123.457 | -67.2099 | -56.2474 |
| nSyb-GS>Tll RNAi 34329 | Tll #2 | 16 | -109.078125 | -76.703125 | -32.375 | -207 | -153.906 | -53.0938 |
| nSyb-GS>Hr96 RNAi 44395 | Hr96 #1 | 30 | -88.89409722 | -93.936111 | 5.042014 | -202.696 | -130.455 | -72.241 |
| nSyb-GS>Usp RNAi 27258 | **Usp #1** | 12 | -88.5 | -5.7916667 | -82.7083 | -209.875 | -181.229 | -28.6458 |
| nSyb-GS>Hr78 RNAi 44393 | Hr78 #1 | 28 | -72.90345982 | -66.556362 | -6.3471 | -139.589 | -73.5385 | -66.0502 |
| nSyb-GS>Ftz-f1 RNAi 27659 | **Ftz-f1 #1** | 10 | -66.409375 | -81.428125 | 15.01875 | -189.2 | -129.892 | -59.3083 |
| nSyb-GS>EcR RNAi 37058 | **EcR #1** | 41 | -64.88378339 | 35.2905052 | -100.174 | -186.678 | -71.3674 | -115.311 |
| nSyb-GS>Hr51 RNAi 39032 | Hr51 #1 | 26 | -46.2948718 | -40.956731 | -5.33814 | -157.457 | -106.372 | -51.0849 |
| nSyb-GS>Tll RNAi 27242 | Tll #1 | 32 | -41.25 | -30.677083 | -10.5729 | -244.021 | -126.177 | -117.844 |
| nSyb-GS>Hr83 RNAi 44397 | Hr83 #1 | 38 | -21.03371711 | -49.833607 | 28.79989 | -65.9433 | -23.7496 | -42.1937 |
| nSyb-GS>Err RNAi 44391 | Err #1 | 23 | -8.710824276 | -95.732111 | 87.02129 | -156.793 | -105.282 | -51.5104 |

E-Gal4 columns are for the subtracted mean values between experimental groups and their Gal4 control groups, and E-UAS columns are for the subtracted mean values between experimental groups and their UAS RNAi control groups. TD, total sleep. RD, daytime sleep. RN, nighttime sleep. N is the number of alive flies for the experimental group. Fly lines bold in the RNA lines column are ecdysone responsive genes.

**Table S2.**
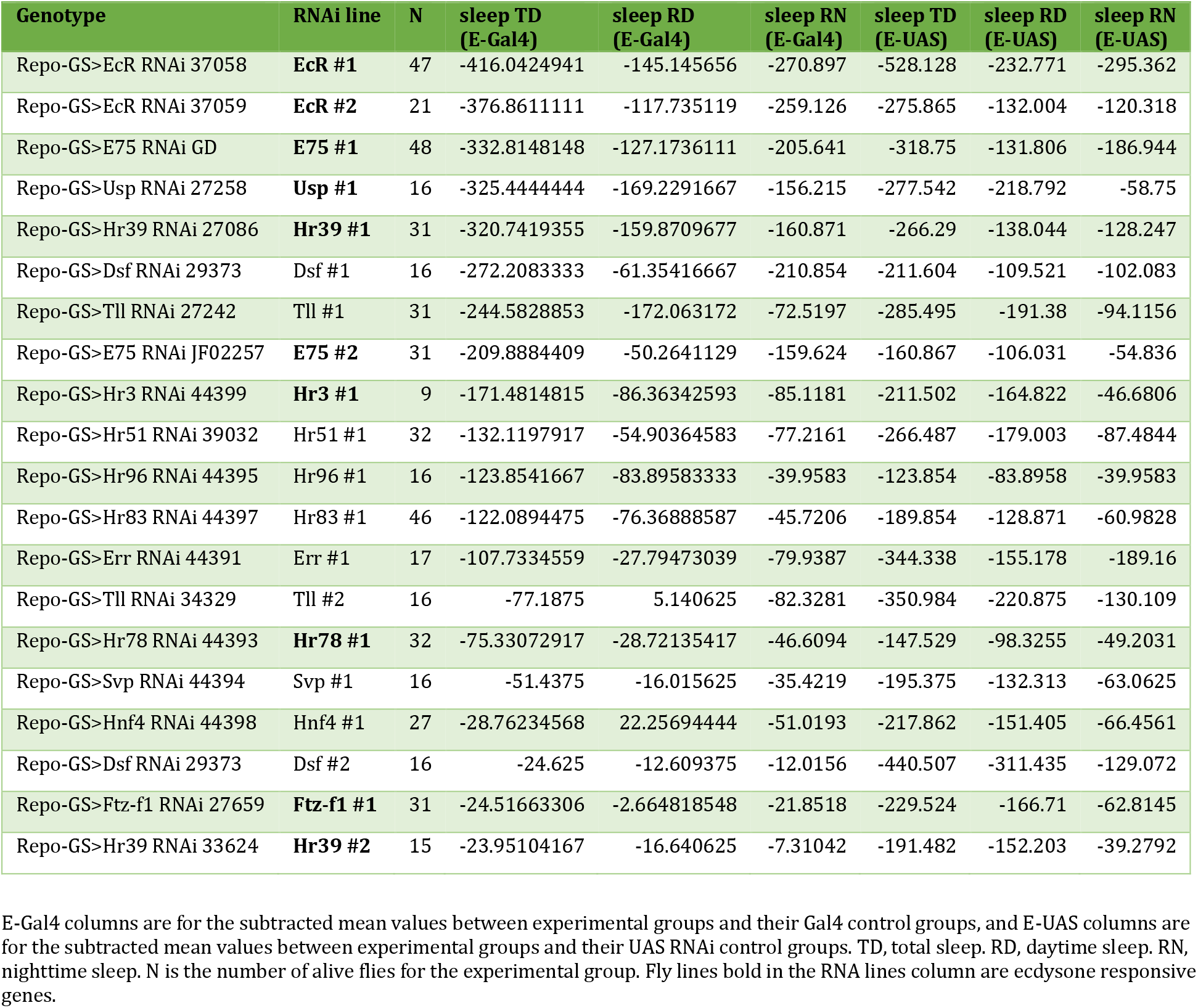
Pan-glia screening of NHRs in adult *Drosophila*

| Genotype | RNAi line | N | sleep TD<br>(E-Gal4) | sleep RD<br>(E-Gal4) | sleep RN<br>(E-Gal4) | sleep TD<br>(E-UAS) | sleep RD<br>(E-UAS) | sleep RN<br>(E-UAS) |
| --- | --- | --- | --- | --- | --- | --- | --- | --- |
| Repo-GS>EcR RNAi 37058 | **EcR #1** | 47 | -416.0424941 | -145.145656 | -270.897 | -528.128 | -232.771 | -295.362 |
| Repo-GS>EcR RNAi 37059 | **EcR #2** | 21 | -376.8611111 | -117.735119 | -259.126 | -275.865 | -132.004 | -120.318 |
| Repo-GS>E75 RNAi GD | **E75 #1** | 48 | -332.8148148 | -127.1736111 | -205.641 | -318.75 | -131.806 | -186.944 |
| Repo-GS>Usp RNAi 27258 | **Usp #1** | 16 | -325.4444444 | -169.2291667 | -156.215 | -277.542 | -218.792 | -58.75 |
| Repo-GS>Hr39 RNAi 27086 | **Hr39 #1** | 31 | -320.7419355 | -159.8709677 | -160.871 | -266.29 | -138.044 | -128.247 |
| Repo-GS>Dsf RNAi 29373 | Dsf #1 | 16 | -272.2083333 | -61.35416667 | -210.854 | -211.604 | -109.521 | -102.083 |
| Repo-GS>Tll RNAi 27242 | Tll #1 | 31 | -244.5828853 | -172.063172 | -72.5197 | -285.495 | -191.38 | -94.1156 |
| Repo-GS>E75 RNAi JF02257 | **E75 #2** | 31 | -209.8884409 | -50.2641129 | -159.624 | -160.867 | -106.031 | -54.836 |
| Repo-GS>Hr3 RNAi 44399 | **Hr3 #1** | 9 | -171.4814815 | -86.36342593 | -85.1181 | -211.502 | -164.822 | -46.6806 |
| Repo-GS>Hr51 RNAi 39032 | Hr51 #1 | 32 | -132.1197917 | -54.90364583 | -77.2161 | -266.487 | -179.003 | -87.4844 |
| Repo-GS>Hr96 RNAi 44395 | Hr96 #1 | 16 | -123.8541667 | -83.89583333 | -39.9583 | -123.854 | -83.8958 | -39.9583 |
| Repo-GS>Hr83 RNAi 44397 | Hr83 #1 | 46 | -122.0894475 | -76.36888587 | -45.7206 | -189.854 | -128.871 | -60.9828 |
| Repo-GS>Err RNAi 44391 | Err #1 | 17 | -107.7334559 | -27.79473039 | -79.9387 | -344.338 | -155.178 | -189.16 |
| Repo-GS>Tll RNAi 34329 | Tll #2 | 16 | -77.1875 | 5.140625 | -82.3281 | -350.984 | -220.875 | -130.109 |
| Repo-GS>Hr78 RNAi 44393 | **Hr78 #1** | 32 | -75.33072917 | -28.72135417 | -46.6094 | -147.529 | -98.3255 | -49.2031 |
| Repo-GS>Svp RNAi 44394 | Svp #1 | 16 | -51.4375 | -16.015625 | -35.4219 | -195.375 | -132.313 | -63.0625 |
| Repo-GS>Hnf4 RNAi 44398 | Hnf4 #1 | 27 | -28.76234568 | 22.25694444 | -51.0193 | -217.862 | -151.405 | -66.4561 |
| Repo-GS>Dsf RNAi 29373 | Dsf #2 | 16 | -24.625 | -12.609375 | -12.0156 | -440.507 | -311.435 | -129.072 |
| Repo-GS>Ftz-f1 RNAi 27659 | **Ftz-f1 #1** | 31 | -24.51663306 | -2.664818548 | -21.8518 | -229.524 | -166.71 | -62.8145 |
| Repo-GS>Hr39 RNAi 33624 | **Hr39 #2** | 15 | -23.95104167 | -16.640625 | -7.31042 | -191.482 | -152.203 | -39.2792 |

E-Gal4 columns are for the subtracted mean values between experimental groups and their Gal4 control groups, and E-UAS columns are for the subtracted mean values between experimental groups and their UAS RNAi control groups. TD, total sleep. RD, daytime sleep. RN, nighttime sleep. N is the number of alive flies for the experimental group. Fly lines bold in the RNA lines column are ecdysone responsive genes.

## STAR⋆Methods

### Key resource table

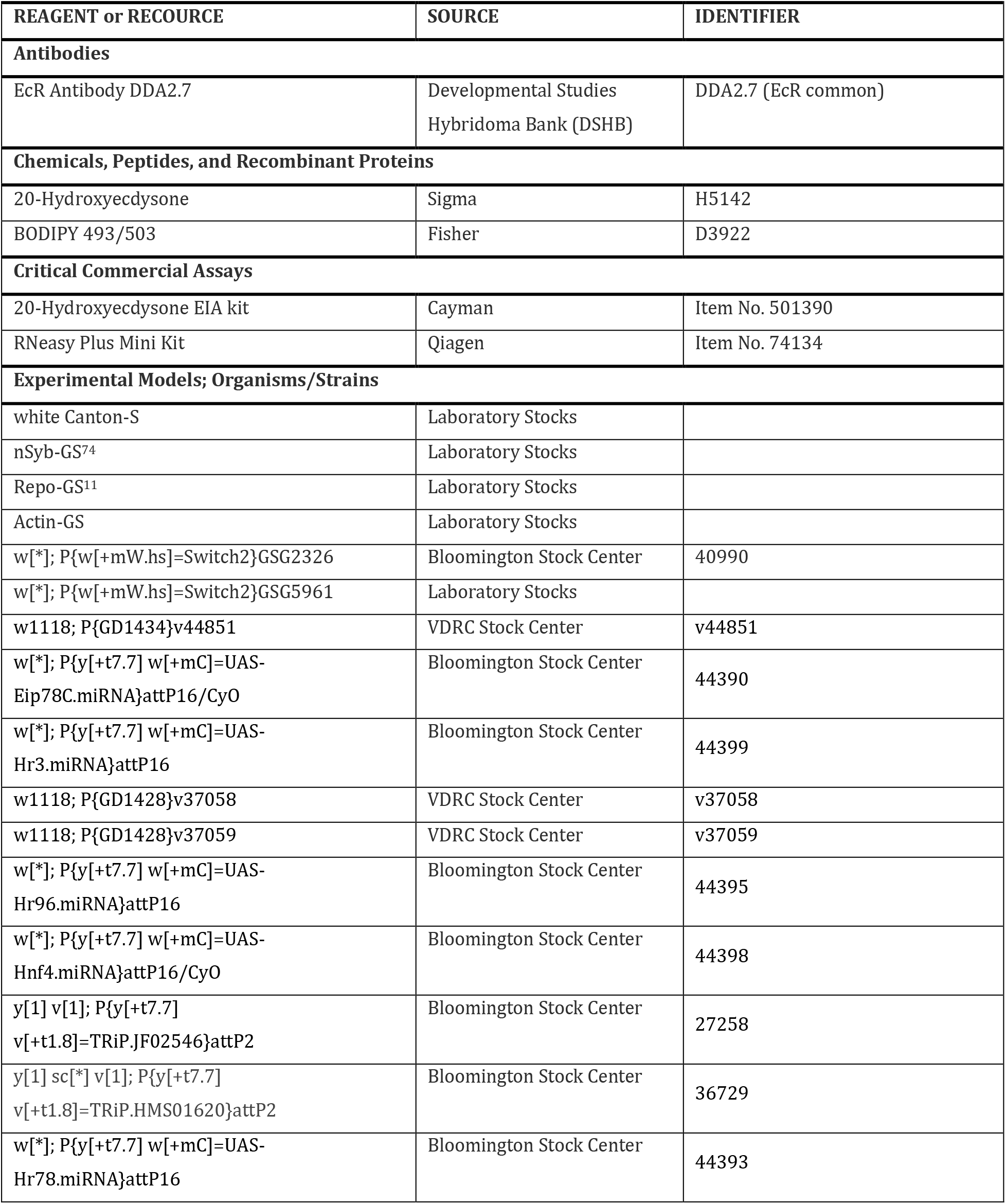
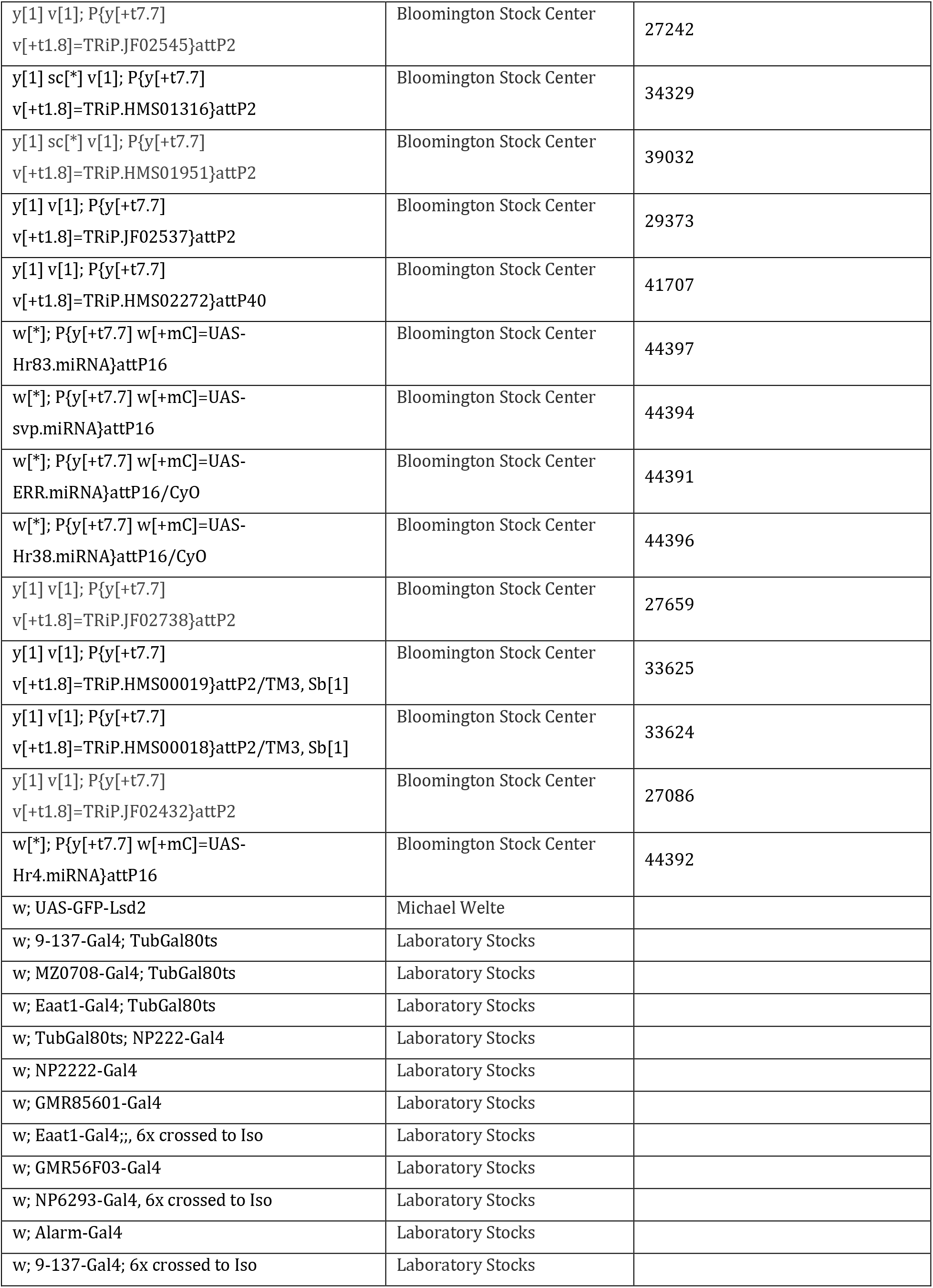
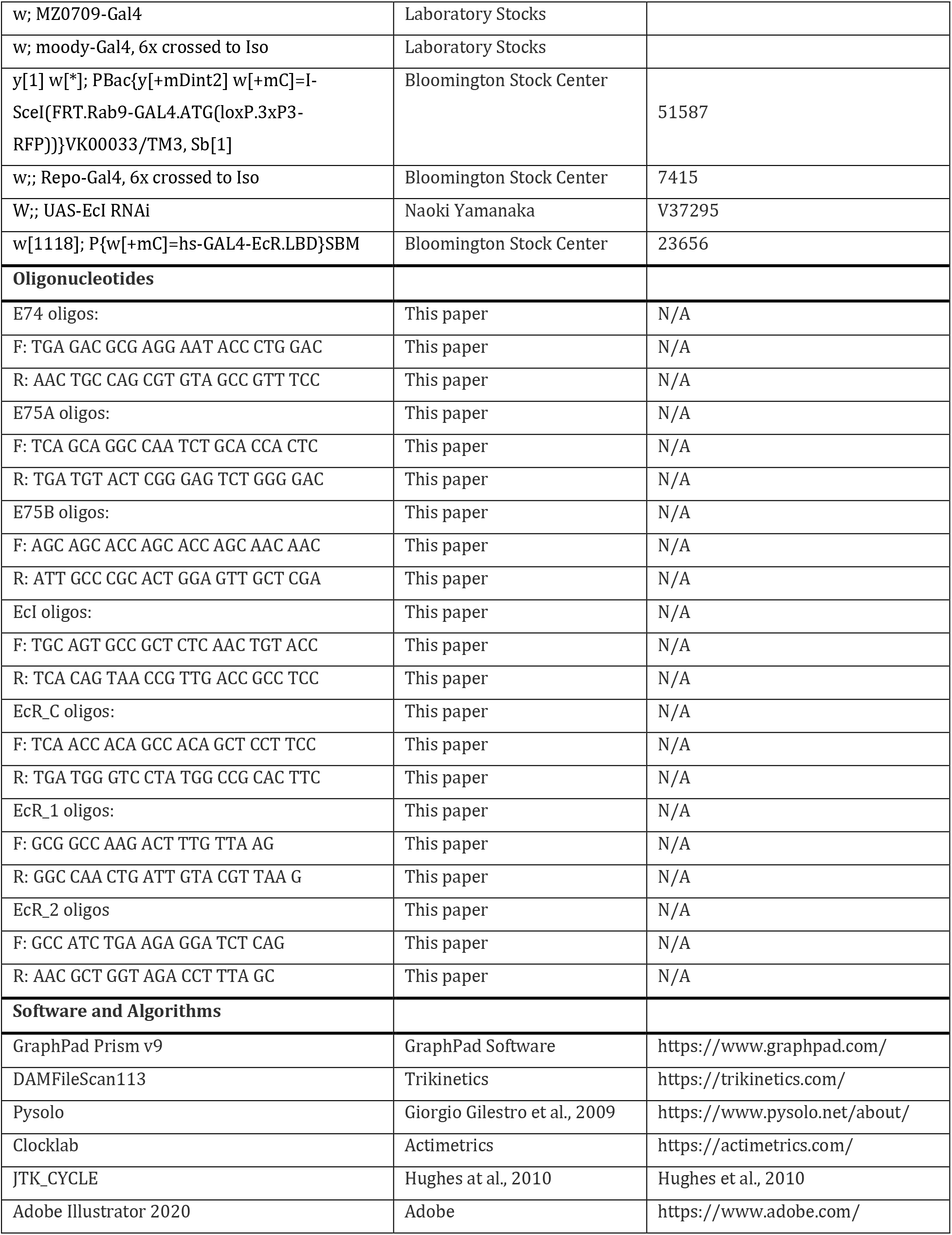

### Contact for reagent and resource sharing

Further information and requests for resources and reagents should be directed to lead contact Amita Sehgal.

### Fly stock and maintenance

*Drosophila* stocks were obtained from our lab stock, Bloomington *Drosophila* stock center (BDSC), or the Vienna *Drosophila* Resource Center (VDRC). The white-CantonS (*w*CS) strain was used as wild-type unless specified. The genotype information of the flies used in each experiment is listed in the key resource table. Strains used from the lab stocks include nSyb-GeneSwitch, Repo-GeneSwitch, all the sub-glial lines, and four sub-glial tubGal80ts lines. nSyb-GeneSwitch, Repo-GeneSwitch experiments were conducted with UAS-Dcr2 to promote RNAi efficiency. EcR RNAi lines are from the lab stock originally purchased from VDRC. The Lsd-2 flies were gifts from Dr. Michael Welts lab, and EcI lines were gifts from Dr. Naoki Yamanaka lab.

### Behavior measurement in *Drosophila*

Flies were raised on cornmeal-molasses medium under 12: 12 h light: dark cycle at 25°C unless specified otherwise. 12 female virgins and four male flies were usually crossed together to generate different genotype flies. Parents flies are cleared after seven days, and F1 progenies are collected after twelve days. On day17, 5-7 days old female flies are loaded into locomotor tubes for behavior tests as previously described (Davla et al., 2020). Locomotor tubes are 60 mm glass tubes vaxed with 2% agar with 5% sucrose as fly food on one side, and the yarn is put on the other side to restrain the behavior of flies inside the glass tubes. For GeneSwitch experiments, 0.5mM RU-486 (mifepristone) was added to the fly food to activate the GeneSwitch. Three constitutive days’ data are used for sleep analysis by Pysolo (https://www.pysolo.net/) (Gilestro and Cirelli, 2009), and seven constitutive days’ data under constant dark are used for circadian rhythm analysis by ClockLab (https://actimetrics.com/products/clocklab/).

### Ecdysone treatment, immunohistochemistry, and imaging

Ecdysone arrived with powder and was dissolved in ethanol. Then it was mixed with 2% agar with 5% sucrose to make the different doses of ecdysone tubes, and the same amount of ethanol as ecdysone solution was used to make the vehicle control tubes. 5–7-day old flies are loaded into locomotor tubes for ecdysone treatment and behavior recording to verify that ecdysone promotes sleep in ecdysone-treated flies.

BODIPY493 was used for brain lipid droplet staining. Flies are loaded into locomotor tubes to get 0.5 mM ecdysone treatment for 12 hours or 36 hours. Both control and ecdysone-treated flies are then dissected in the PBS, fixed in the 4% PFA for 20 mins, and washed three times with PBS + 0.3% triton (PBT). Then brains will be left in PBT at 4 degrees overnight and transferred to 1ug/ml BODIPY493 in PBT for 20 mins. Later they are mounted for imaging.

Brains are imaged with the oil-immersion 40x of confocal microscope with a resolution of 1024X1024. Raw images are processed with FIJI ImageJ. The first step is to remove non-lipid trash and exclude artificial signals or signals outside the brain area. The second step is to quantify the lipid droplet (LD) count, area, and total brain tissue area. Later, the lipid droplets count will be normalized by the brain tissue area, and lipid droplet size is calculated by LD area divided by total count. Quantifications were conducted by using ImageJ Macro.

### Ecdysone assay

20E measurement is performed with the 20-hydroxyecdysone EIA kit (Cayman, item No. 501390). 5 Whole flies, 10 fly heads, or 30 brains are dissected and homogenized in the 70% ethanol, and protein levels are measured by the BCA assay for normalization. The homogenates were then dried by the evaporator, redissolved in the kit’s buffer, and then measured based on the manufacturer’s protocol accordingly.

### Statistical analysis

GraphPad Prism was used for all statistical tests. Data were tested for normality using the D’Agostino-Pearson test and Shapiro-Wilk test. Normally distributed data were then tested with an unpaired parametric student t-test for two independent groups and a one-way ANOVA analysis with Turkey post-hoc test for multiple independent groups. Non-normally distributed data, such as sleep bout numbers, which are usually non-normally distributed, were analyzed with a nonparametric test like the Mann-Whitney test for two samples and the Kruskal-Wallis test with Dunn’s multiple comparisons test for three samples or above. Statistic tests used for each experiment are indicated in the figure legends, and data are presented as means and standard error of the mean (SEM).

## Data availability

All data generated and analyzed for this study are included in the manuscript and supporting files.

## Supplemental information

Supplemental Information includes six figures and two tables.

